# FLASH: A next-generation CRISPR diagnostic for multiplexed detection of antimicrobial resistance sequences

**DOI:** 10.1101/426338

**Authors:** Jenai Quan, Charles Langelier, Alison Kuchta, Joshua Batson, Noam Teyssier, Amy Lyden, Saharai Caldera, Aaron McGeever, Boris Dimitrov, Ryan King, Jordan Wilheim, Maxwell Murphy, Lara Pesce Ares, Katherine A. Travisano, Rene Sit, Roberto Amato, Davis R. Mumbengegwi, Jennifer L. Smith, Adam Bennett, Roly Gosling, Peter M. Mourani, Carolyn S. Calfee, Norma F. Neff, Eric D. Chow, Peter S. Kim, Bryan Greenhouse, Joseph L. DeRisi, Emily D. Crawford

## Abstract

The growing prevalence of deadly microbes with resistance to previously life-saving drug therapies is a dire threat to human health. Detection of low abundance pathogen sequences remains a challenge for metagenomic Next Generation Sequencing (NGS). We introduce FLASH (Finding Low Abundance Sequences by Hybridization), a next-generation CRISPR/Cas9 diagnostic method that takes advantage of the efficiency, specificity and flexibility of Cas9 to enrich for a programmed set of sequences. FLASH-NGS achieves up to 5 orders of magnitude of enrichment and sub-attomolar gene detection with minimal background. We provide an open-source software tool (FLASHit) for guide RNA design. Here we applied it to detection of antimicrobial resistance genes in respiratory fluid and dried blood spots, but FLASH-NGS is applicable to all areas that rely on multiplex PCR.

## INTRODUCTION

Emerging drug resistant pathogens represent one of the most significant threats to human health. Drug resistant infections currently claim 700,000 lives per year and are predicted to cause 10 million deaths annually by 2050(1). Antimicrobial susceptibility information is crucial to implement targeted and effective therapeutic interventions, but is often unobtainable due to the need to first isolate a pathogen in culture(2), a process that can require days to months depending on the organism, and has low success rates in the setting of prior antibiotic use(3, 4). Ongoing and emerging drug resistance is a central challenge for malaria and other parasitic diseases as well(5, 6). To limit the spread and impact of anti-malarial drug resistance, real-time surveillance of resistance patterns is essential. Current methods include *in vivo* efficacy studies from patient samples, *in vitro* phenotypic resistance assays and PCR-based detection of gene mutations associated with drug resistance(7). While rapid genotyping offers many advantages over organism viability studies, in areas of high disease transmission it is confounded by the presence of co-infections where a low-abundance strain may contain clinically and epidemiologically relevant sequences essential for assessing transmission patterns(8).

Metagenomic Next Generation Sequencing (mNGS) has proven invaluable for detecting pathogens in clinical samples(9–11); however, the key issue of antimicrobial resistance (AMR) detection is not easily addressed by mNGS alone. While interrogation of antibiotic resistance genes is readily achievable from cultured isolates, it is often not possible from direct clinical specimens due to low target abundance and high background derived from the host. Thus, detection of low abundance targets is a central challenge in clinical diagnostics, and a solution would have universal relevance across medical disciplines. Methods combining multiplex PCR with NGS, such as 16S rRNA gene profiling and AmpliSeq(12) provide effective enrichment but are hampered by cost, scalability and inflexibility when new targets are discovered. Other approaches that rely on probe-based hybrid capture suffer from high off-target rates, long incubation times, and expensive reagents.

The exquisite and programmable specificity of CRISPR systems has inspired many novel uses of their enzymes beyond genome engineering since the characterization of *S. pyogenes* Cas9 in 2012(13). The SHERLOCK(14, 15) and DETECTR(16, 17) methods take advantage of Cas13 and Cas12a to detect limited sets of pathogen sequences with attomolar sensitivity in clinical samples. Our group recently demonstrated that recombinant Cas9 coupled with multiplexed sets of guide RNAs can be used for precision depletion of unwanted background sequences(18). We have now built on that work and developed FLASH (Finding Low Abundance Sequences by Hybridization).

This novel NGS targeted enrichment system has direct applicability to the challenge of AMR detection, among other applications. The FLASH technique uses a set of Cas9 guide RNAs designed to cleave sequences of interest into fragments appropriately sized for Illumina sequencing (Figure 1). Input genomic DNA or cDNA is first blocked by phosphatase treatment and then digested with Cas9 complexed to this set of guide RNAs. The resulting cleavage products are thus made competent for ligation of universal sequencing adapters. With the ensuing amplification, the targeted sequences are enriched over background and made ready for binding to the sequencing flow cell. This method goes beyond other CRISPR-based diagnostic tools in that it enables high levels of multiplexing (thousands of targets) and is reinforced by the precision and sequence identity confirmation that is inherent in a traditional NGS readout. We highlight two uses of FLASH-NGS in the realm of drug-resistant infections: the burden of antimicrobial resistance genes in pneumonia-causing gram-positive bacteria and drug resistance in the malaria parasite *Plasmodium falciparum*.

**Figure 1.**
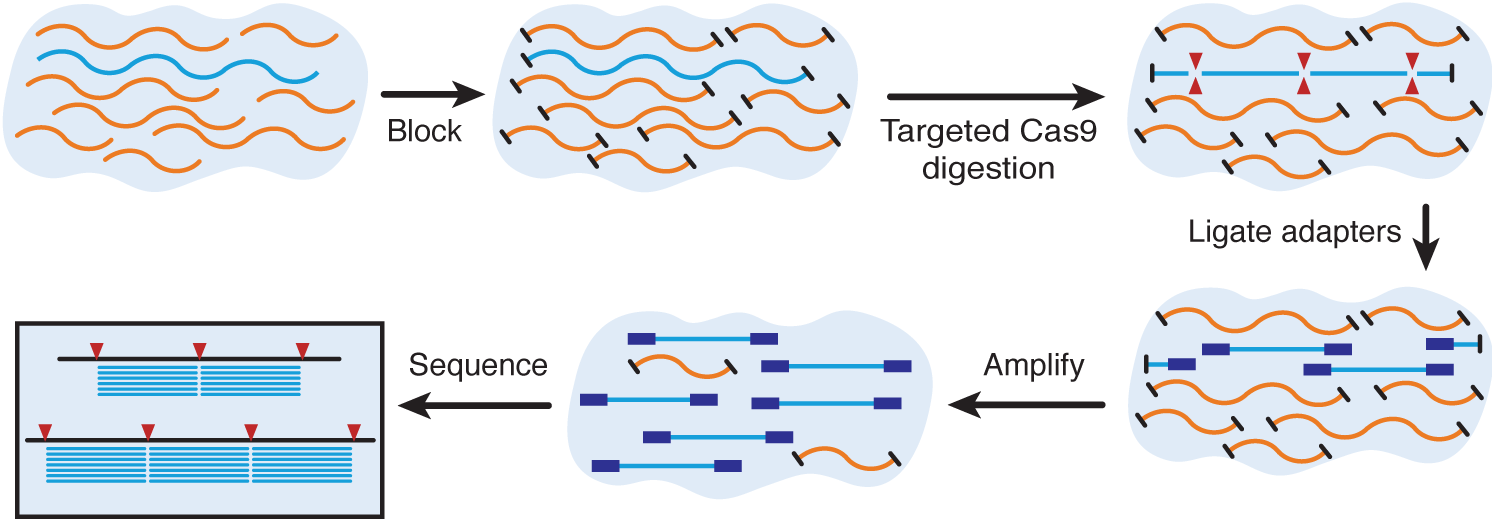
Overview of the FLASH method. Genomic DNA or cDNA is first blocked with phosphatase treatment and then digested with Cas9 complexed to a set of guide RNAs targeting genes of interest. Ligation of sequencing adaptors, amplification and sequencing follows.

## MATERIALS AND METHODS

To choose optimal guide RNA targets for FLASH, we developed a flexible computational tool called FLASHit. Given a set of target genes, this tool first defines targetable 20-mer Cas9 sites, applying certain exclusion criteria (see Supplementary Methods). It then takes advantage of homology between genes to design a relatively small set of guide RNAs that provides a relatively high sequence coverage (Figure S1). While a single FLASH-derived fragment is sometimes sufficient to uniquely identify an AMR gene, we designed guides to cut each gene into multiple fragments, both to increase the probability of detection in the case of a single nonfunctional guide RNA or unanticipated SNP and to maximize the ability to detect both known and unknown sequence variants. This was achieved by solving a mixed integer program with the objective of maximizing the number of inter-guide inserts of optimal Illumina NGS length (200-300 bp) covering each gene while minimizing the number of guide RNAs (see Supplementary Methods and github documentation). FLASHit is freely available at github.com/czbiohub/flash.

### Bacterial AMR FLASH

In order to construct a limited pilot set of guide RNAs that would be compatible with a more comprehensive future set, FLASHit was first used to design a set targeting the full collection of 3,624 clinically relevant AMR-related genes derived from the CARD(19) and ResFinder(20) databases, merging exact duplicates (Table S1). This set contained 5,513 guide RNAs (Table S2). For pilot experiments, a subset of these sequences was chosen to target 127 clinically relevant AMR genes present in *Staphylococcus aureus* and other gram-positive bacteria (Table S3), including 118 acquired resistance genes and 9 chromosomal genes capable of carrying drug resistance-conferring point mutations (indicated in Table S4). The latter mainly represent highly conserved genes and serve two functions: to determine the presence of a given bacterial species (even if no acquired resistance genes are present) and to identify resistance point mutations. This set contained 532 target sequences (indicated as “staph” in Table S2), with the majority of genes containing at least four target sites (Figure S1). DNA templates for producing crRNAs (CRISPR RNAs) for each target were synthesized, transcribed separately, then purified and pooled.

For the cultured isolate experiments, DNA was isolated from six clinical *S. aureus* isolates and sequenced in triplicate with traditional NGS (NEBNext Ultra II FS DNA-Seq kit) and FLASH-NGS using the pilot guide RNA set described above, to a sequencing depth greater than 0.5M reads for each replicate. For a detailed FLASH-NGS protocol, see Supplementary Methods. Briefly, 5’ phosphate groups were enzymatically cleaved using rAPid alkaline phosphatase which was subsequently deactivated with sodium orthovanadate. The dephosphorylated DNA was added to a master mix containing the CRISPR/Cas9 ribonucleoprotein complex and incubated at 37°C for 2 hours. The Cas9 was deactivated with proteinase K and removed with a SPRI bead purification Samples were dA-tailed and adapter-ligated using the NEBNext Ultra II reagents and protocols. Following two SPRI bead purifications to remove adapter dimer, samples were indexed with 22 cycles of PCR using dual unique TruSeq i5/i7 barcode primers. Individual libraries were pooled and size selected for fragments between 250-600; pools that were too low in concentration were subsequently amplified using up to 5 cycles of PCR. All libraries were sequenced on Illumina MiSeq or NextSeq instruments. Data were demultiplexed, quality filtered with PriceSeqFilter(21), aligned to the 127 targeted *S. aureus* genes using Bowtie 2(22), and tallied up with custom python scripts. Unless otherwise noted, all reported data represents reads per million (rpM) averaged across triplicates. Filtering and alignment results, including no-template controls, are provided in Table S5.

In the direct clinical sample experiments, for each sample 25 ng of DNA or cDNA was subjected to NGS and FLASH-NGS following the protocol outlined above and described in detail in Supplementary Methods. Data were aligned to our pilot set of 127 targeted gram-positive genes using Bowtie 2(22). Filtering and alignment results, including no-template controls, are provided in Table S5.

### Plasmodium FLASH

Six *P. falciparum* genomic loci with drug-resistance associations (D-01 to D-06), 25 with high population diversity (P-01 to P-25) and 17 microsatellite sites (M-01 to M-17) were selected for FLASH-NGS (Figure S6). A single guide RNA target site was chosen from each side of each locus such that PE150 sequencing of the resulting insert would yield coverage of all SNPs of interest (Table S8). In order to best simulate clinically relevant samples, dried blood spots (DBSs) representing three different mixtures of the culture adapted *P. falciparum* strains U659, HB3 and D10 were prepared by spotting 20 µL of blood containing 10,000 parasites/µL onto filter paper. DNA was subsequently extracted and amplified by selective whole genome amplification (sWGA) using custom primers(23). One hundred nanograms of DNA from each sample was subjected to FLASH-NGS in the manner described above, using the *P. falciparum* guide RNA set. This was repeated for three independent sWGA reactions from each of the three different mixtures, as well as each of the three strains alone. We also sequenced sWGA-amplified mixed-strain blood spots without FLASH as a control. Each dataset consisted of at least 2M PE150 reads. Reads were aligned to the Pf3D7 genome (PlasmoDB version 28(24)) using Bowtie 2(22) and further analysis was done using SAMtools(25), BEDTools(26) and custom python and R scripts. Filtering and alignment results, including no-template controls, are provided in Table S5.

## RESULTS AND DISCUSSION

### Assessing FLASH using cultured bacterial isolates

FLASH performance for AMR gene sequence identification was first evaluated in the context of cultured bacterial isolates. DNA from six clinical *S. aureus* isolates was sequenced in triplicate with traditional NGS and FLASH-NGS using the pilot guide RNA set described above. As expected, all nine chromosomal genes were recovered in all six isolates by NGS. Each isolate also contained between zero and four acquired resistance genes. For all isolates, every gene identified by NGS was also identified by FLASH-NGS above a threshold of 1,000 rpM (Figure 2A). In a single case, one false positive gene was identified above this threshold with FLASH-NGS: ErmC at 6,461 rpM in isolate 6. We believe this to be the result of cross-contamination from another isolate, either during library prep or on the sequencer, as ErmC was one of the two most abundant genes identified across all FLASH-NGS isolates (Figure 2A and Table S5). This result highlights the need for caution when multiplexing samples using extremely sensitive amplification techniques such as FLASH. To summarize, in this experiment on cultured isolate DNA (where the ground truth was given by unbiased NGS), we observed zero false negatives and one false positive when considering 127 target genes across 6 samples.

**Figure 2.**
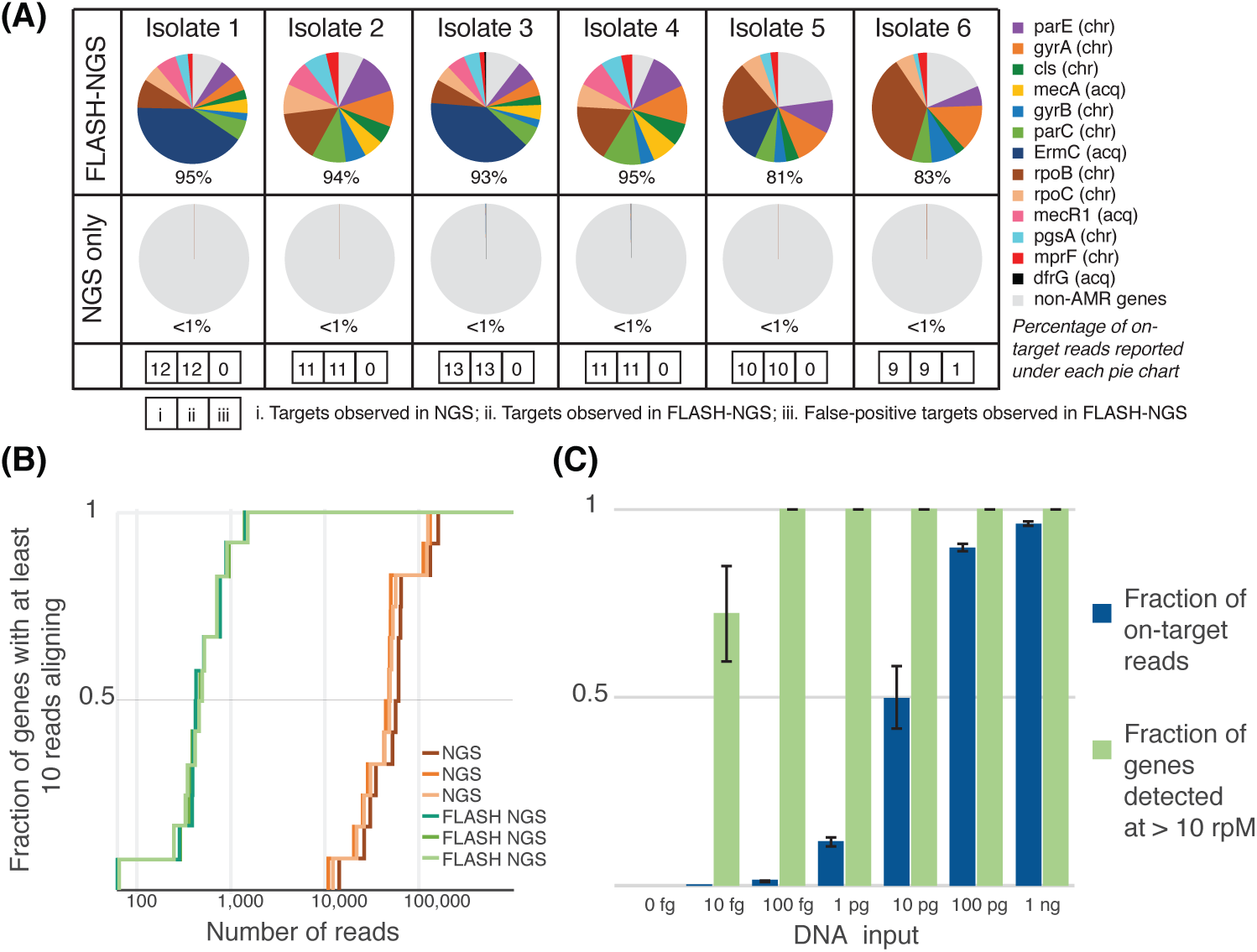
Results of FLASH on cultured isolates. (A) Proportion of targeted genes detected in FLASH-NGS and NGS libraries of six *S. aureus* isolates. Average of 3 replicates. chr: chromosomal gene; acq: acquired gene. Numbers in boxes represent, in order for each isolate: i. number of targets present (based on NGS); ii. number of expected targets observed in FLASH-NGS; iii. number of false-positive targets observed in FLASH-NGS (false positive not depicted in pie chart). (B) For isolate 1 with FLASH, less than 2,000 sequencing reads were needed to achieve coverage of each targeted gene by at least 10 reads. Over 100,000 reads were needed to achieve the same coverage with NGS alone. (C) The fraction of targeted reads relative to background decreases substantially below 100 pg of DNA input; however, with as little as 100 fg input (approximately 35 copies of the *S. aureus* genome), the full set of targeted genes was detected at 10 rpM or greater. Bars and error bars represent mean and standard deviation of three replicates.

The FLASH-NGS results were consistent with phenotypic testing, with the exception of one ciprofloxacin resistant isolate, and three instances of non-*ermC* clindamycin resistance – these exceptions were to be expected given that our pilot target set did not cover the full spectrum of ciprofloxacin or clindamycin resistance elements (Table S6). On average, 90.1% of reads mapped to target genes in FLASH samples, compared to 0.3% mapping to these genes with NGS alone. This represented a 293-fold increase in average rpM of targeted genes (Figure 2A). For each FLASH-NGS sample, a sequencing depth between 500 and 5,000 reads was sufficient to recover 10 or more reads per gene for 100% of targeted genes. For NGS alone, at least 100-fold higher sequencing depth was required to achieve this minimal threshold (Figure 2B and Figure S2).

Several parameters may affect FLASH-NGS performance, including the complexity of the guide RNA set, the amount of input DNA, and the amount of Cas9 protein. The effect of guide RNA pool complexity was tested using an extended set of 2,226 guides (transcribed as a single pool) (Figure S3, Table S2). On-target performance remained comparable, with an observed 90.6% recovery rate. To determine the limits of input nucleic acid for FLASH profiling, the mass of DNA from isolate 1 was progressively lowered to as little as 10 femtogram (Figure 2C). The fraction of FLASH-derived reads dropped below 50% at 10 pg (replaced mostly by reads corresponding to *E. coli* derived from the Cas9 preparation itself; see Table S5). Despite fewer on-target reads, all targeted genes were still covered by at least 10 rpM at 100 fg (approximately 35 *S. aureus* genome copies in 30 µL, or 1.9 aM), and over half of them were covered at 10 fg (0.19 aM). The amount of input Cas9 protein had little effect on the enrichment of target sequences down to 0.4pmol, which corresponds to approximately 50 copies of each Cas9-guide RNA complex per *S. aureus* genome copy. This represents a materials cost of < $1 US when using commercially available Cas9 and guide RNAs transcribed from crRNA templates in a pool (Figure S4).

We examined the *S. aureus* isolate data to understand why a small number of guide RNAs yielded few or no reads (Table S7). Whole genome sequence data was used to determine the sequences at the target locations for all *S. aureus* isolates. Considering every target site in every gene present in each of the six isolates, there were a total of 622 target sequences present in this set of experiments (approximately 100 for each isolate). A total of 94.4% (587 of 622) target sites were readily detectable in FLASH-NGS experiments, using an arbitrary but conservative detection threshold of 1 rpM across the three replicates. Of the remaining 35 target sites, we noted that 9 (25.7%) harbored mutations within the targeting sequence. This reinforces the need to build in additional redundancy by selecting multiple target sites per gene, a key feature of the FLASHit program. It is unclear exactly why the remaining 26 guide RNAs failed. While numerous tools exist to assess the efficiency of different guide RNA sequences(27, 28), they are mostly concerned with *in vivo* genome editing capabilities, rather than *in vitro* cutting activity. We expect that as more FLASH data is collected, *in vitro* guide failure patterns will emerge.

### FLASH enrichment directly from clinical samples

Culture-based infectious disease diagnostics often fail to provide actionable data due to challenges with growing fastidious and slowly replicating microbes such as mycobacterial and fungal pathogens, and administration of antibiotics prior to sampling. We thus sought to assess the performance of FLASH-NGS for detecting AMR gene targets directly from four patient samples with culture-confirmed drug resistant infections (Table S6), including from both DNA and RNA preparations. Each was subjected to NGS and FLASH-NGS following the protocol outlined above and described in detail in Supplementary Methods. Data were aligned to our pilot set of 127 targeted gram-positive genes using Bowtie 2(22). Filtering and alignment results, including no-template controls, are provided in Table S5. Figure 3 shows reads per million for each targeted gene in each clinical sample detected with NGS alone and FLASH-NGS. All data represent averages of three to six experiments. In the context of patient samples, where the microbial component is a minority of the nucleic acid, the proportion of on-target sequences was lower than for cultured isolates (Figure S5). However, the mean enrichment over NGS was > 5,000-fold (range 563-fold to 13,244-fold per sample), and in many cases FLASH-NGS detected genes that were unobservable with NGS alone, as described below. Due to the difficulty of assessing true positivity in the context of these clinical samples, it is difficult to determine sensitivity and specificity of the FLASH method in this context. We note that studies currently underway on broader sets of clinical samples will address this question more directly.

**Figure 3.**
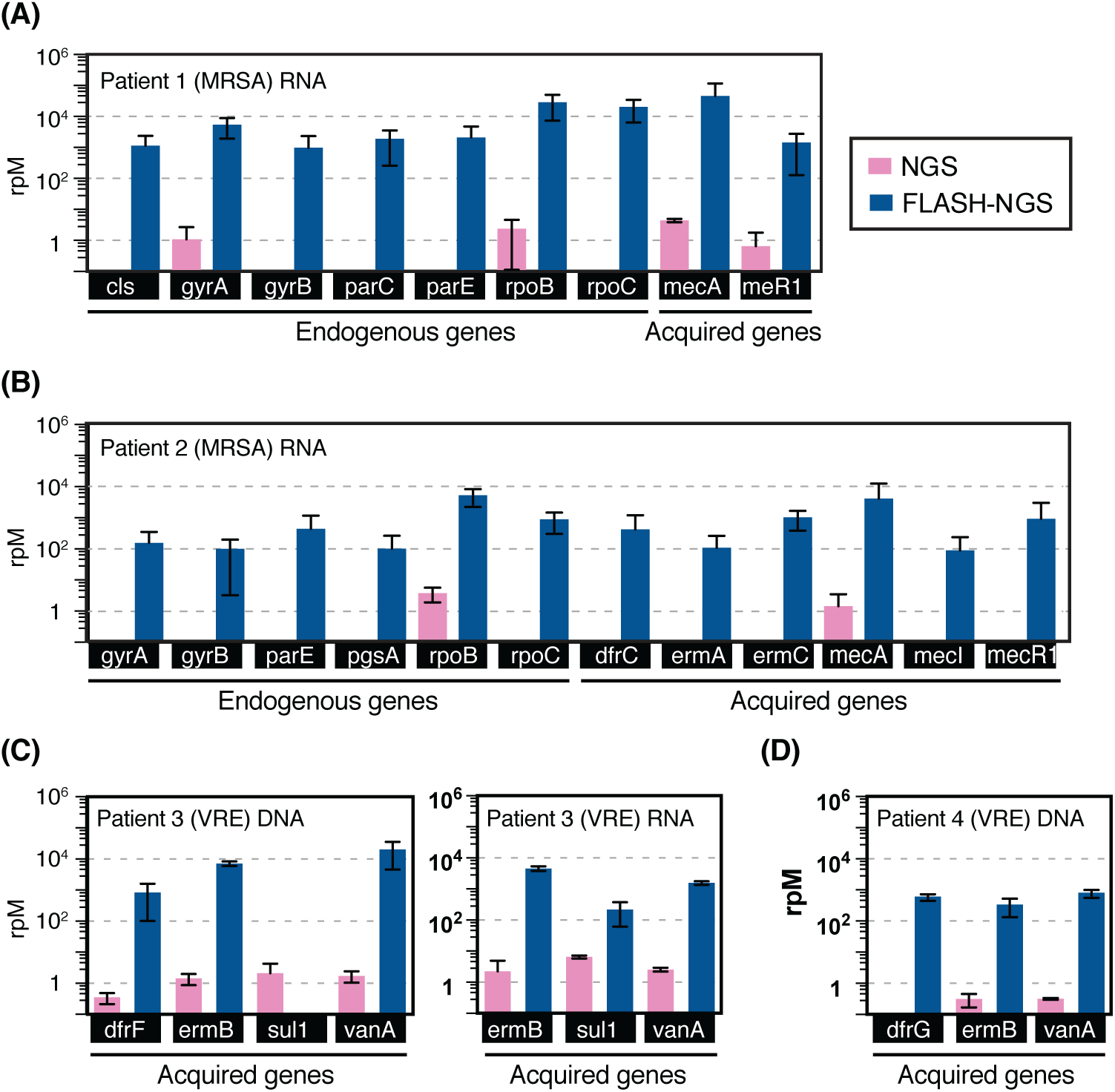
Results of FLASH on respiratory samples. Number of reads aligning to targeted genes in NGS and FLASH-NGS sequencing experiments on respiratory fluid samples from (A) patient 1, (B) patient 2, (C) patient 3, and (D) patient 4. Average of 3 or 6 replicates. Bars and error bars represent mean and standard deviation of three to six replicates (see Table S5).

Patient 1 (Figure 3A) was hospitalized for culture-positive methicillin-resistant *Staphylococcus aureus* (MRSA) pneumonia. FLASH-NGS of RNA extracted from mini-bronchial alveolar lavage (mBAL) identified the *mecA* gene, which explains the methicillin resistance observed in the *S. aureus* isolated from this patient (Table S6), at over 20,000 rpM. Seven *S. aureus* chromosomal genes were also detected (*cls*, *gyrA*, *gyrB*, *parC*, *parE*, *rpoB* and *rpoC*). Patient 2 was admitted with fatal pneumonia that was culture positive for MRSA, *Citrobacter* and *Pseudomonas*. FLASH-NGS performed on RNA extracted from tracheal aspirate (TA) detected six *S. aureus* chromosomal genes (*gyrA*, *gyrB*, *parE*, *pgsA*, *rpoB* and *rpoC*) plus acquired genes conferring resistance to trimethoprim-sulfamethoxazole (TMP-SMZ) (*dfrC*), macrolides (*ermA* and *ermC*) and methicillin (*mecA* and its regulators *mecI* and *mecR1*). Given the polymicrobial nature of this patient’s infection, it was not possible to say with certainty whether the acquired resistance genes originated with the phenotypically multidrug resistant *S. aureus* (Table S6) or from another species.

Patient 3 was admitted for a lower respiratory tract infection. Vancomycin-resistant *Enterococcus faecium* (VRE) was identified by culture. FLASH-NGS of DNA from mBAL fluid identified the *vanA* gene, which confers vancomycin resistance. Resistance to macrolides and TMP-SMZ are widespread in *Enterococci* and thus identification of *ermB* (macrolide resistance), and of *dfrF* (TMP-SMZ resistance) was not surprising. With FLASH-NGS of RNA from the same sample, *dfrF* was not detected (Figure 3C), likely indicating this gene was not being expressed. We note that *sul1* was detected in patient 3 DNA by NGS but not FLASH-NGS. Examination of all read pairs aligning to this gene in this patient revealed that they exclusively mapped to a 250 bp region containing only a single FLASH target site. This does not preclude FLASH enrichment: in RNA, thousands of read pairs were derived from this site on one side and nonspecific cleavage on the other side. Why this was not observed in the DNA samples is unknown. Overall, the patient 3 results demonstrate the utility of testing both nucleic acid types. RNA profiling provides information on what genes are active, and relatively high RNA expression levels may enhance the probability of detection. DNA profiling may identify relevant genotypes even in the absence of expression, results that will at times be clinically relevant given the inducible nature of some AMR mechanisms

Patient 4 (Figure 3D) was admitted for VRE bacteremia and also found to have vancomycin-resistant *Enterococcus faecium*; FLASH-NGS of DNA from this patient’s TA identified *vanA* (vancomycin resistance) as well as *ermB* (macrolide resistance) and *dfrG* (TMP-SMZ resistance).

In addition to acquired resistance elements, antimicrobial resistance can also be conferred by single point mutations in chromosomal genes. Notably, FLASH-NGS can recover SNP data simultaneously with presence/absence data for acquired resistance genes located on mobile genetic elements. For example, RNA from patient 2 shows wildtype sequence for all but one rifampicin resistance SNP location in the *rpoB* gene. At position 481, where wildtype was histidine, we found a mixture of 33% wildtype and 77% H481Y mutation out of a total of over 20,000 reads. H841Y *rpoB* has been described as rifampicin resistant (29). We also found 97.3% of the over 500 reads mapping to *gyrA* position 84 in this patient represented the S84L mutation, which explains the ciprofloxacin resistance observed in the S. aureus isolated from this patient (Table S6)(19). The genetic mixtures may indicate a coinfection with more than one strain of *S. aureus* in this patient, or evolving resistance mutations.

### FLASH-NGS for *Plasmodium falciparum* drug resistance and strain diversity

The challenge of multiplex drug resistance detection is not limited to prokaryotes. We adapted FLASH to identify malaria strain variants in the context of mixed infections, an important challenge for malaria control and elimination efforts and a modality that could be extended to many areas of epidemiology. Six *P. falciparum* genomic loci with drug-resistance associations (D-01 to D-06), 25 with high population diversity (P-01 to P-25) and 17 microsatellite sites (M-01 to M-17) were selected for FLASH-NGS (Figure S6) and a single guide RNA target site was chosen from each side of each locus as described in Methods. DBSs representing three different mixtures of the culture-adapted *P. falciparum* strains U659, HB3 and D10 were prepared and then sequenced with NGS and FLASH-NGS as described in Methods. On average we observed 85.6% on-target reads with FLASH-NGS on these samples, compared with less than 0.02% on-target without FLASH (Figure S7). For the triple strain samples, the 31 D and P windows targeted were enriched from two to more than five orders of magnitude over traditional NGS when averaging all experiments from all strain mixtures (n = 9, three replicates each of three different strain mixes) and considering only read pairs in which all haplotype-defining SNPs in a particular window were sequenced (Figure 4A).

**Figure 4.**
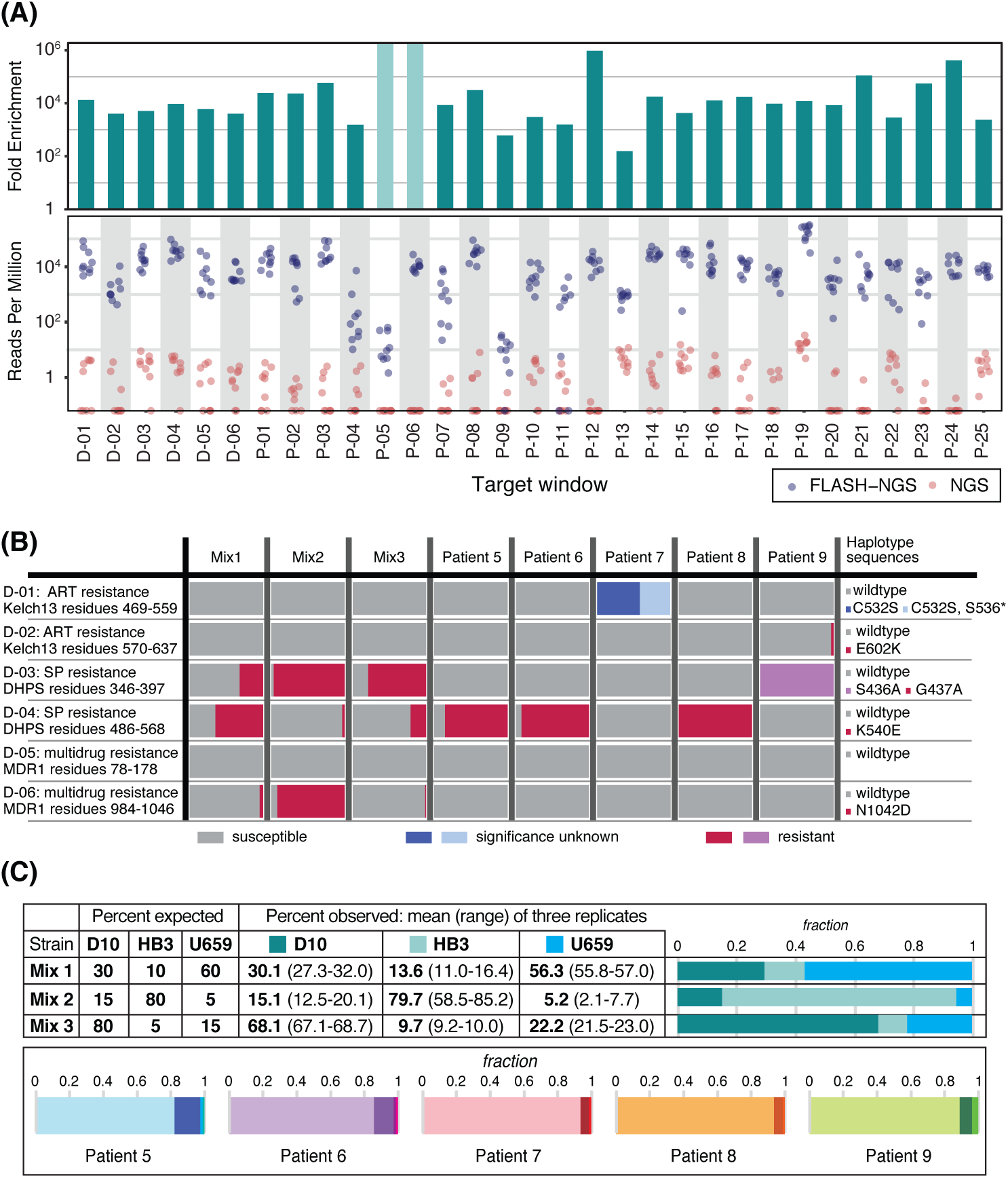
Results of FLASH on dried blood spots. (A) Dried blood spots (DBSs) from malaria lab strains were sequenced using either NGS or FLASH-NGS. Reads per million is plotted for each window for nine FLASH-NGS samples (three strain mixtures, each in triplicate, blue) and nine equivalent NGS samples (pink). Top panel indicates fold enrichment (average FLASH-NGS rpM divided by average NGS rpM). Light green bars in the upper panel represent windows for which no haplotype-determining read pairs were found. (B) Both lab strain mixtures and patient samples were evaluated for the presence of drug resistant haplotypes. Bars indicate mean of three replicates. (C) Target windows with sequences that distinguish the lab strains D10, HB3 and U659 were used to estimate strain ratios in the three mixtures. For the clinical DBS samples, the number of sequencing reads attributable to different haplotypes at each window was determined using SeekDeep. The average proportion of haplotypes at each of the maximum-haplotype containing windows is depicted. Bars indicate mean of three replicates.

We also applied FLASH-NGS with the same guide RNA set to DBSs from patients in the Zambezi region of Namibia (patients 5-9). DBSs were selected for having a parasite density greater than 10,000/µL (determined by qPCR, see Supplementary Methods). These were extracted, amplified with sWGA and subjected to FLASH-NGS in triplicate. The six targeted drug resistance windows represented loci in the genes *kelch-13*, *dhps* and *mdr1* (Figure 4B). Using FLASH-NGS, at least 2500 sequencing reads were obtained for each window for each sample (it should be noted, however, that in this pilot guide RNA set only two windows were targeted for each gene, so additional non-targeted mutations could have been missed). Mutations in *kelch-13* confer resistance to artemisinin. No *kelch-13* mutations were detected in the lab strains. Patient 7 had a *kelch-13* C532S mutation at 100%; this has been observed before but the significance is unknown(30). In addition, 41.4% of reads in this patient had a stop codon mutation at position 536. Patient 9 had an E602K mutation present in 2.8% of reads, suggesting a low frequency of artemisinin resistance. Mutations in the *dhps* gene conferring resistance to the antimalarial Sulfadoxine/Pyrimethamine (specifically the resistance-conferring K540E mutation) are relevant for public health in Africa as their prevalence is used by the WHO to determine intermittent presumptive therapy (IPT) regimens. We detected the resistant G437A haplotype at variable levels (along with the susceptible wildtype haplotype) in all three lab strain mixtures. Four of the patients had mutations in this region as well: patients 5, 6 and 8 had the resistant K540E mutation at 85-100%, and patient 9 had the resistant S436A mutation at 100%. This result was expected in this population(31). Finally, we targeted the multidrug resistance locus *mdr1*. The lab strain mixtures showed variable levels of the N1042D resistance mutation, but only wildtype *mdr1* sequences were detected in the patient samples.

*P. falciparum* haplotypes derived from FLASH-NGS experiments on lab strain mixtures were analyzed to determine whether minor strains were detectable and whether strain ratios could be accurately recovered. Twenty-one of the 48 windows uniquely distinguished all three strains from each other (Figure S8). In all three mixtures, we reliably detected all three strains. To estimate the haplotype ratios from these data, we averaged the percentages across these windows (Figure 4C). Figure S8 depicts the percentages of each strain represented in each window (dots) along with the median and interquartile range across all windows (boxes). While any individual window was an imprecise estimate of haplotype ratio, averaging across the 21 windows converged on an accurate estimate for each of the three replicates of each of the three strain mixtures.

For the clinical samples, the number of variable haplotypes was determined for each window using SeekDeep(32), and is depicted in Figures 4C and S9. Patients 5 and 6 have two windows each for which four unique haplotypes were identified, and patients 7, 8 and 9 have at least 4 windows each for which three unique haplotypes were identified. Averaging strain percentages across only these maximum haplotype windows indicated that the primary haplotype comprised between 82% and 94% of parasites, with the 2-3 additional haplotypes making up the remainder (Figure 4C). Notably, none of these patients shared a complete set of identical primary haplotype sequences, suggesting that five different strains accounted for the primary infections in these five individuals (although some windows did share identical sequences) (Figure S9).

## CONCLUSION

In conclusion, we have developed a targeted sequencing method that is fast, inexpensive, has high multiplexing capacity, and is nimble enough to target virtually any sequence of interest without optimization. It is the efficiency, specificity and programmability of the CRISPR/Cas9 system that allows this functionality. Highly multiplexed detection of antimicrobial resistance genes in patient samples is an important use case for FLASH-NGS; however, it is by no means the only application area for this technique. Detection of mutations in cancer, rare mosaic allele detection, targeted transcriptomics from clinical samples, enrichment of microbiome components from complex mixtures, and recovery of targeted transcripts from single cell sequencing libraries are among other possible applications that may be explored in future work.

## AVAILABILITY

Microbial sequence data are available on SRA in BioProject ID PRJNA493248. Code is available on github.com/czbiohub/flash.

## ACKNOWLEDGEMENT

We are extremely thankful to Dr. Sofonias Tessema for assistance with the malaria work. We also thank Anna Sellas, Gorica Margulis, Anna Chen and Jennifer Mann for general laboratory help. We appreciate Drs. Michael Wilson, Wei Gu, Matthew Zinter, Brian O’Donovan, Christine Sheridan and Greg Fedewa for contributing to the conception of the project, and Katrina Kalantar, James Wang, Dr. James Webber and Dr. David Dynerman for advice and assistance with data analysis. Finally, we thank Drs. Stephen Quake and Cristina Tato for assistance with manuscript review. This publication uses data from the MalariaGEN *P. falciparum* Community Project (www.malariagen.net/projects/p-falciparum-community-project). MalariaGEN’s genome sequencing was performed by the Wellcome Sanger Institute and the Community Projects is coordinated by the MalariaGEN Resource Centre with funding from the Wellcome Trust (098051, 090770).

## SUPPLEMENTARY DATA

### Materials and Methods

#### Generation of DNA from *Staphylococcus aureus* isolates

Bacterial cultures were inoculated following standard clinical protocols for tracheal aspirate. Phenotypic antibiotic resistance was obtained using the Vitek2 system (Biomerieux, Marcy-l’Étoile, France) (Table S6). DNA from cultured isolates was extracted using either the Zymo Quick DNA extraction kit (Zymo Research, Irvine, CA) or the Qiagen AllPrep RNA/DNA kit (Qiagen, Hilden, Germany) according to manufacturer’s instructions.

#### Generation of DNA and cDNA from direct clinical respiratory specimens

Excess tracheal aspirate (patients 1 and 4) or mini-bronchial alveolar lavage (patient 3) specimens were collected from patients enrolled in a study examining acute respiratory disease in critically ill adults according to University of California San Francisco Institutional Review Board protocol 10-02701. Tracheal aspirate specimen from patient 2 was collected from a patient enrolled in a study investigating factors predisposing to ventilator associated pneumonia in critically ill children according to University of Colorado Institutional Review Board protocol 14-1530. DNA and RNA was extracted using the Qiagen AllPrep RNA/DNA kit on the QIAcube as described in (1). cDNA was generated from extracted RNA using Nugen Ovation v2 SPIA amplification kit (NuGen, San Carlos, CA, USA). Phenotypic antibiotic resistance was obtained using the Vitek2 system (Table S6).

#### *Plasmodium falciparum* dried blood spot extractions and selective whole genome amplification

Individual asexual *P. falciparum* lab strains D10, HB3 and U659 were grown in human donor red blood cells (RBCs) at 2% hematocrit (percentage of isolated and cleaned RBCs in total culture volume) in RPMI 1640 media with 2mM L-Glutamine, 25 mM HEPES, 2 g/L sodium bicarbonate, 5 g/L AlbuMAX II Lipid-Rich BSA (Life Technologies, Carlsbad, CA, USA), 0.1 mM hypoxanthine, 50 mg/L gentamicin. Cultures were grown at 37°C, 5% O_2_, and 5% CO_2_. RBCs were collected from human donors under UCSF IRB number 10-3852. Parasitemia (percentage of parasite-infected RBCs out of total RBCs) and synchronization of the parasite culture was routinely monitored.

To simulate clinical dried blood spot (DBS) samples, cultured parasites were spotted onto Whatman filter paper with a parasite density of 10,000 parasites/spot as determined by % parasitemia and % hematocrit. This was done for each of the three lab strains (D10, HB3 and U659) individually, and also for cultures consisting of the three strains mixed at three different known percentages (Fig. 4C). Thus, a total of 6 DBS sample types were used, each in triplicate. Genomic DNA was isolated from the dried blood spots using the QiaAMP blood DNA mini kit from Qiagen.

Five samples were collected from symptomatic malaria cases in 2016 from the Zambezi region of Namibia as described (Tessema et al., in preparation). Briefly, samples were spotted on 3MM Whatman filter paper, dried and stored at −20°C until processing. Six mm hole punches from dried blood spots were extracted using the saponin Chelex method (2) and parasite density was quantified using varATS ultra-sensitive qPCR (3). Hole punching and extraction was done in triplicate for each of the five DBSs.

All DBS-extracted DNA samples underwent selective whole genome amplification (sWGA), an isothermal amplification technique that allows for preferential enrichment of the malaria genome over the human genome, as described previously with some modifications (4, 5). Briefly, a 50 µL sWGA PCR reaction consisting of a final concentration of 1X NEB phi29 reaction buffer, 1X Bovine Serum Albumin, 2.5 µM primer set 6A (4), 2.5 µM of primer set 10A (5), 2 mM dNTP mix with 70% AT and 30% GC ratio, 30 units of NEB phi29 DNA Polymerase, plus 20 µL of DNA directly from the Qiagen or Chelex extraction. Cycling conditions are as follows: 35°C x 5 min, 34°C x 10 min, 33°C x 15 min, 32°C x 20 min, 31°C x 30 min, 30°C x 16 hours, 65°C x 15 min, 6°C hold. Reactions were quantified by high sensitivity DNA Qubit (Thermo Fisher Scientific, Waltham, MA, USA).

#### Preparation of CRISPR/Cas9

The CRISPR/Cas9 (Clustered Regularly Interspersed Short Palindromic Repeats/CRISPR associated protein 9) protein was generated as described in (6) From N to C terminus, the construct consisted of the following components: 6X HIS tag, maltose binding protein (MBP), S. pyogenes Cas9, 2X SV40 nuclear localization site (NLS), mCherry, 1X SV40 NLS.

Briefly, the Cas9 vector was expressed in BL21-CodonPlus (DE3)-RIL competent cells (Agilent, Santa Clara, CA, USA) for three hours at 16°C, after which cultures were centrifuged and frozen. Thawed cell pellets were later resuspended in lysis buffer (50 mM sodium phosphate pH 6.5, 350 mM NaCl, 10% glycerol, 1 mM TCEP) supplemented with protease inhibitors and microfluidized. The soluble fraction was purified on a heparin column on the GE AKTA Pure system, then concentrated down and further fractionated by size exclusion chromatography. The resulting pooled fractions were concentrated and stored at −80°C in 50% glycerol.

#### Guide RNA design, antimicrobial resistance genes

We began with all antimicrobial resistance genes listed in the CARD and ResFinder databases, removing exact duplicates. Using FLASHit, we determined the set of all possible guide RNAs (100,931) with the following exclusion criteria: (a) no homopolymers of length greater than 5; (b) no runs of 2-base repeats greater than 3; (c) no internal hairpins; and (d) GC content between 25% and 75%. Reasoning that a guide that can bind a human sequence could be sequestered away from its pathogen target by the highly abundant human DNA in a metagenomic sample, FLASHit removes from consideration any guide that matches a human sequence with zero mismatches in the five PAM proximal bases (the seed region (7, 8)), with one or fewer mismatches in the 10 most PAM proximal bases, and with 2 or fewer mismatches in the full 20-mer. FLASHit also removes guides that match a sequence in the common *E. coli* expression strain BL21, used to produce the Cas9, to avoid false positives resulting from residual DNA bound to the enzyme. FLASHit was then used to define an optimized set of 5,513 guides from this set. The optimization method uses a mixed integer program (9) and is described in the github documentation. Characteristics of the resulting guide RNA set are depicted in Fig. S1.

#### Guide RNA design, *P. falciparum* drug resistance windows, population diversity windows, and microsatellite windows

Areas of significant diversity across the malaria genome were identified by using custom R scripts to search for 250 bp regions with multiple single nucleotide polymorphisms (SNPs) having global minor allele frequencies of > 0.1. These targets have epidemiological diversity between multiple populations. Twenty-five of them were selected and are referred to as population diversity windows (P). Seventeen microsatellites regions (M) were also included. Six windows located at three key malaria drug resistance genes of interest (*kelch-13*, *dhps* and *mdr1*) were also selected, and are referred to as drug resistance windows (D). Potential target RNA binding sites were identified for each window. Pairs of guide RNAs surrounding each insert were chosen by hand based on coverage with PE150 sequencing (i.e. window size less than 300bp) and absence of knowns SNPs at the guide RNA target site. All guide RNA target sites had GC content between 5% and 50% and none contained a homopolymer greater than 9 bases in length or a series of greater than 5 dinucleotide repeats or greater than 3 trinucleotide repeats. Insert sizes ranged from 156 bp to 338 bp, with an average of 245 bp. Key SNP positions within each window were identified from MalariaGEN with a minor allele frequency (MAF) >= 0.001 in Africa (https://www.malariagen.net/projects/p-falciparum-community-project). Chromosomal locations of target windows are indicated in Fig. S6. Windows, guide RNA targets, and SNPs are indicated in Table S8.

#### Guide RNA preparation

DNA oligonucleotide templates for CRISPR RNA (crRNA) and trans-activating CRISPR RNA (tracrRNA) were purchased from Integrated DNA Technologies (IDT, Coralville, IA, USA). crRNA template sequences were as follows (T7 RNA polymerase site is underlined):

5’<u>TAATACGACTCACTATAG</u>NNNNNNNNNNNNNNNNNNNNGTTTTAGAGCTATGCTGTTTTG 3’

where the 20 Ns represent the 20 nucleotide target region. tracrRNA template sequence was as follows:

5’<u>TAATACGACTCACTATAG</u>GACAGCATAGCAAGTTAAAATAAGGCTAGTCCGTTATCAACTT GAAAAAGTGGCACCGAGTCGGTGCTTTTT3’

crRNAs were transcribed individually from their DNA template using T7 RNA polymerase for 2 hours at 37°C. Each reaction contained the following components: 100 ng crRNA DNA template or 8ug tracrRNA DNA template, T7 buffer (final concentrations 40 mM Tris pH 8.0, 20 mM MgCl2, 5 mM DTT, and 2 mM spermidine), NTPs (1mM each ATP, CTP, UTP, GTP), and 10 ng/µL T7 enzyme. Guide RNAs were purified using SPRI (Solid Phase Reversible Immobilization) magnetic beads. Individual crRNAs were pooled and stored as 80 µM single-use aliquots at −80°C. Immediately prior to use, pooled crRNAs were annealed with tracrRNA at an equimolar ratio to form 40 µM dual-guide RNA.

For the expanded set of 2,226 antimicrobial resistance (AMR) guide RNAs (Fig. S3), all guides were transcribed in a single pool.

#### Preparation of standard Next Generation Sequencing (NGS) Libraries

RNA libraries from patient respiratory fluid: Five µL of RNA from each sample was reverse transcribed with the NuGEN Ovation v2 SPIA kit. The concentration of cDNA was quantified via high sensitivity DNA Qubit. End repair of 25 ng of cDNA was done as described in the NEBNext Ultra II protocol (New England Biolabs, Ipswitch, MA, USA).

DNA libraries from patient respiratory fluid: After extraction, DNA was quantified via high sensitivity DNA Qubit. Twenty-five nanograms of DNA was fragmented for 12.5 minutes at 37°C and end-repaired at 65°C using the NEBNext Ultra II FS DNA protocol.

DNA libraries from bacterial cultured isolates: Samples of between 10 fg and 25 ng of total DNA were fragmented for 5 minutes at 37°C and end-repaired at 65°C using the NEBNext Ultra II FS DNA protocol.

DNA libraries from sWGA of malaria DBS: One hundred nanograms of sWGA-amplified DNA was end-repaired as described in the NEBNext Ultra II protocol.

All libraries: Following end-repair, adaptor ligation at a 1:100 adaptor dilution was performed as described in the NEBNext Ultra II protocol. A SPRI bead purification was done at a sample:bead volume ratio of 1:1 and samples were eluted in 15 µL. The purified ligated products were indexed with 9-18 cycles of PCR using NEB Q5 polymerase and dual unique TruSeq i5/i7 barcode primers. A SPRI bead purification was done at a sample:bead volume ratio of 1:1. Individual libraries were analyzed using the High Sensitivity DNA Bioanalyzer kit and pooled based on library shape and according to the concentration of DNA within the range of 250 to 600 base pairs. Pooled samples were size selected for fragments between 250-600 bp using the BluePippin 2% agarose gel cassette (Sage Science, Beverly, MA, USA). The BluePippin product was SPRI bead purified at a sample:bead volume ratio of 1:1.4 and eluted in 15 µL of water. Pooled library quality was assessed on the Bioanalzyer (Agilent technologies, Santa Clara, CA). Pools that were too low in concentration were amplified using up to 5 cycles of KAPA HiFi HotStart Real-time PCR (Kapa Biosystems, Roche, Basel, Switzerland) according to the manufacturer’s protocol and cycling conditions. Amplified samples were SPRI bead purified using a sample:bead volume ratio of 1:1.4 and eluted in 15 µL. Pooled libraries were quantified using High Sensitivity DNA Qubit. Final library quality was assessed using on the Bioanalyzer and additional SPRI bead purifications were performed if adapter dimer was present. Final libraries were quantified using qPCR.

#### Preparation of FLASH-NGS Libraries

The 5’ phosphate groups of 25 ng DNA or cDNA from cultured isolates or respiratory fluid, or 100 ng of sWGA from malaria DBSs, were enzymatically cleaved using rAPid alkaline phosphatase (Sigma Aldrich, St. Louis, MO, USA) for 30 mins at 37°C according to the manufacturer’s instructions. The phosphatase enzyme was deactivated with one unit of sodium orthovanadate. The dephosphorylated DNA was added to a master mix containing the CRISPR/Cas9 ribonucleoprotein complex. The final mixture was a 30 µL reaction of dephosphorylated DNA, Cas9 buffer (50 mM Tris pH 8.0, 100 mM NaCl, 10 mM MgCl_2_, and 1 mM TCEP), 500 nM Cas9, and 6 µM appropriate dual guide RNAs. The mixture was incubated at 37°C for 2 hours. The Cas9 was deactivated by adding 1 µL of proteinase K and incubating at 37°C for another 15 minutes. A SPRI bead purification at a sample:bead volume ratio of 1:1.7 was conducted and samples were eluted in 50 µL. Samples were dA-tailed using the NEBNext dA-Tailing Module. Adaptor ligation at a 1:100 adaptor dilution was performed as described in the NEBNext Ultra II protocol. Two SPRI bead purifications were done at a sample:bead volume ratio of 1:1 and samples were eluted in 15 µL. The purified ligated products were indexed with 22 cycles of PCR using NEB Q5 polymerase and dual unique TruSeq i5/i7 barcode primers. A SPRI bead purification was done at sample:bead volume ratio of 1:1. Individual libraries were analyzed using the High Sensitivity DNA Bioanalyzer kit and pooled based on library shape and according to the concentration of DNA within the range of 250 to 600 base pairs. Pooled samples were size selected for fragments between 250-600 bp using the BluePippin 2% agarose gel cassette. The BluePippin product was SPRI bead purified at a sample:bead volume ratio of 1:1.4 and eluted in 15 µL. Pooled library quality was assessed on the Bioanalzyer. Pools that were too low in concentration were amplified using up to 5 cycles of KAPA HiFi HotStart Real-time PCR according to the manufacturer’s protocol and cycling conditions. Amplified samples were SPRI bead purified using a sample:bead volume ratio of 1:1.4 and eluted in 15 µL. Pooled libraries were quantified using High Sensitivity DNA Qubit. Final library quality was assessed using on the Bioanalyzer and additional SPRI bead purifications were performed if adapter dimer was present. Final libraries were quantified using qPCR.

#### Sequencing

All libraries were sequenced on Illumina MiSeq or NextSeq instruments (Illumina, San Diego, CA, USA). Raw sequence data for all isolate experiments is available on SRA. Sequence reads mapping to AMR targets or to the *P. falciparum* genome are also available on SRA. Raw read counts, filtered read counts, and number of reads aligning to targets for all experiments are available in Table S5.

#### Cultured S*. aureus* isolate data analysis

Datasets were demultiplexed and then filtered with PriceSeqFilter (10) using the flags *-pair both rqf 85 0.98 -rnf 90*. This removes any read pairs for which either read contains less than 85% of nucleotides with a probability of being called correctly of at least 0.98, or for which either read has less than 90% of nucleotides called (more than 10% Ns). Filtered datasets were aligned to the 127 target genes using Bowtie 2 with the flags *-a -X 1000 --very-sensitive*. The *-a* flag forces all alignments to be considered and reported. The resulting .sam file was filtered using SAMtools (11) with *samtools view -F 256* to retain only the highest-scoring alignment for each read pair. Reads aligning to the 127 target genes were tabulated with custom python scripts. NGS data was used to identify which of the 127 genes was present in each of the six isolates at 10 rpM (reads per million) or greater, and only those genes were included in the analyses. The 127 target genes included two different *parE* sequences from two different strains of *S. aureus*; reads aligning to these two genes were summed to get a *parE* total. These results are presented in Figs. 2, S2, S3 and S4. The additional guide RNAs used in Fig. S3 are marked as “extended” in Table S2.

To determine *E. coli* contamination from the Cas9 prep, filtered datasets were aligned to the *E. coli BL21* genome using Bowtie 2 with the *-very-sensitive-local* flag.

NGS data was used to identify the sequence of each guide RNA target site present in each of the isolates. Only target sites with at least 1 rpM (averaged over the three replicates) having an overlap of at least one base with the site sequence in the NGS isolate data were included in this analysis. Mutations were identified and are indicated in Table S7. For each guide site in each isolate, the alignment files were examined and reads that aligned with a start position within 2 bp of the cut site were identified and tallied as FLASH-derived reads. These results are presented in Table S7.

#### Patient respiratory fluid data analysis

Datasets were demultiplexed and then filtered with PriceSeqFilter using the flags *-pair both -rqf 85 0.98 -rnf 90*. Filtered datasets were aligned to the 127 target genes using Bowtie 2 with the flags *-a -X 1000 --very-sensitive*. The *-a* flag forces all alignments to be considered and reported. The resulting .sam file was filtered with *samtools view -F 256* to retain only the highest-scoring alignment for each read pair. Reads aligning to the 127 target genes were tabulated with custom python scripts. These results are presented in Fig. 3 and S5. Genes were plotted in Fig. 3 only if an average of 100 rpM in FLASH-NGS samples and/or 1 rpM in NGS samples aligned to them. The SNP analysis of the *rpoB* and *gyrA* genes from Patient 1 was done using the variant call feature in Geneious (12).

#### *P. falciparum* data analysis

Enrichment of *P. falciparum* target windows: Datasets were demultiplexed and filtered with PriceSeqFilter (10) using the flags *-pair both -rqf 85 0.98 -rnf 90*. Fig. S7 was generated by using Bowtie 2 to align filtered datasets to Pf3D7 reference sequences of the 48 target windows.

Enrichment of haplotype-determining SNPs in *P. falciparum* target windows: Filtered datasets were also aligned to the Pf3D7 genome (PlasmoDB version 28 (13)) using Bowtie 2 with the flag *--very-sensitive-local*. The resulting .bam files were filtered to contain only properly paired reads with *samtools view -f 0×2,* and then converted to .bedpe files with *bedtools bamtobed -bedpe* (14). These .bedpe files were reduced to .bed files describing the outer spans of each read pair using awk (*gawk ′{ print $1 ″\t″ $2 ″\t″ $6 ″\t″ $7 ″\t″ $8 ″\t″ $9 ″\t″ $10 }′*). A separate .bed file was created containing chromosomal locations encompassing only the key SNP positions (identified by MalariaGEN has having an MAF >= 0.001 in Africa) within each of the 48 target windows (rows denoted “snp_range” in Table S8). Finally, *bedtools intersect* was used to count the number of read pair spans which covered 100% of a target window using the flags *-f 1 -c* for each sample. M (microsatellite) windows were omitted from this analysis. These results are presented in Fig. 4A and represent 9 NGS and 9 FLASH-NGS experiments: three replicates of each of three triple strain mixtures.

Determination of *P. falciparum* haplotype ratios: Single strain NGS datasets from each of the three lab strains was used to determine the sequence of each target window in each strain. Twenty-one of the 48 windows had unique sequences for each of the three strains (P-01, P-03, P-05, P-06, P-07, P-08, P-11, P-12, P-13, P-14, P-15, P-17, P-18, P-21, P-23, P-24, P-25, M-12, M-15, M-17, and M-04). A fasta file containing each sequence for each of these windows was created, and for all NGS and FLASH-NGS experiments, filtered datasets were aligned to this file using Bowtie 2 with the flags *-a -X 1000 --very-sensitive*. The *-a* flag forces all alignments to be considered and reported. The resulting .sam files were filtered with *samtools view -F 256* to retain only the highest-scoring alignment for each read pair. The number of reads aligning to each variant of each target window was tabulated using grep commands. These results are presented in Figs. 4C and S8.

Haplotype analysis of malaria patient samples: Datasets were demultiplexed and then filtered with PriceSeqFilter (10) using the flags *-pair both -rqf 85 0.98 -rnf 90*. Haplotype analysis was carried out using SeekDeep (15) with default parameters. These results are presented in Fig. 4B.

*P. falciparum* drug resistance mutations in lab samples and patient samples: To evaluate drug resistance SNPs, all filtered samples were aligned to the 6 drug resistance windows using Bowtie 2 with the *--very-sensitive-local* flag and analyzed with the variant call feature in Geneious. These results are presented in Fig. 4C.

##### Acknowledgements

This publication uses data from the MalariaGEN *P. falciparum* Community Project (www.malariagen.net/projects/p-falciparum-community-project). MalariaGEN’s genome sequencing was performed by the Wellcome Trust Sanger Institute and the Community Projects is coordinated by the MalariaGEN Resource Centre with funding from the Wellcome Trust (098051, 090770)

---

**Figure S1.**
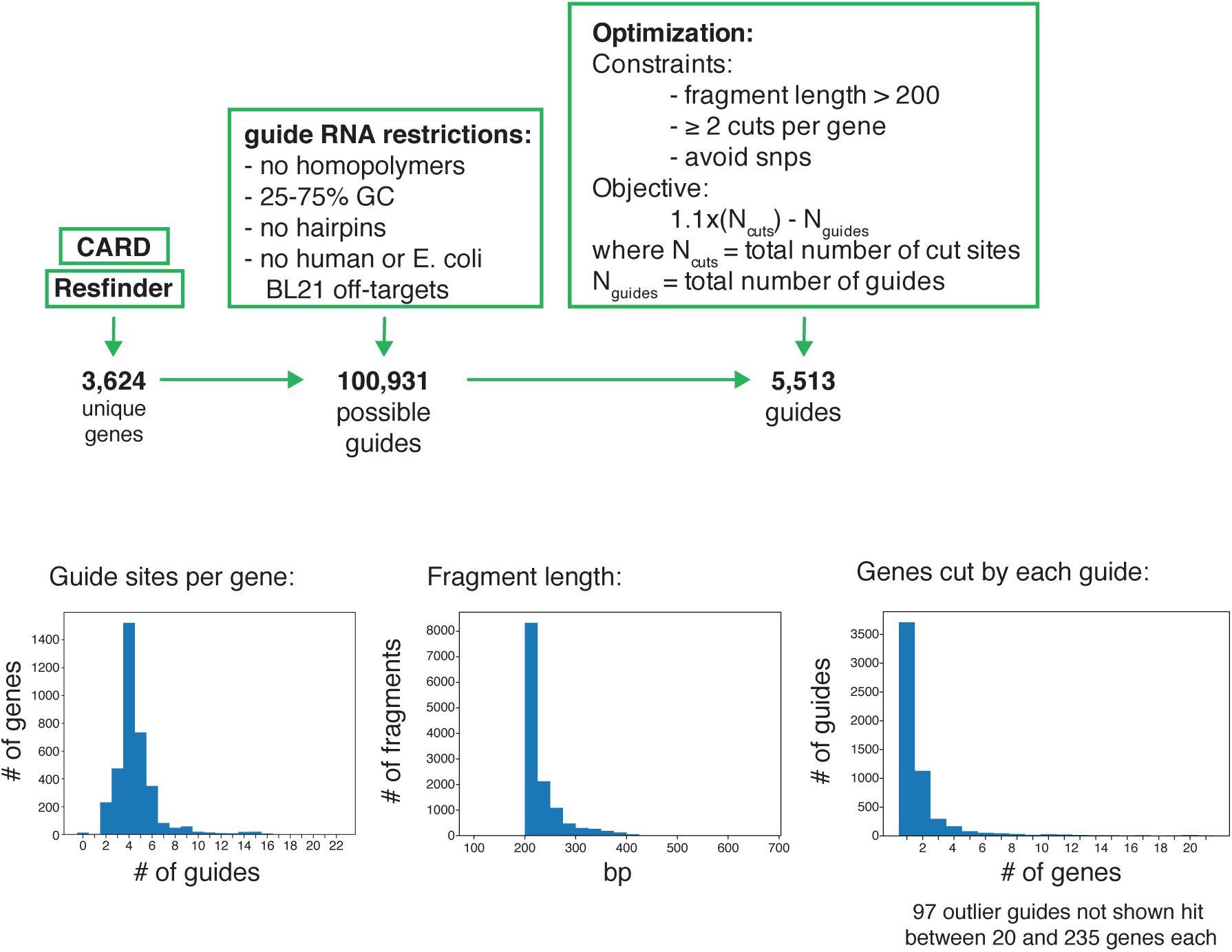
Top: Antimicrobial resistance (AMR) gene guide RNA set design strategy. Bottom: Histograms showing properties of the 5,513 guide RNA set designed to target all known bacterial AMR genes.

**Figure S2.**
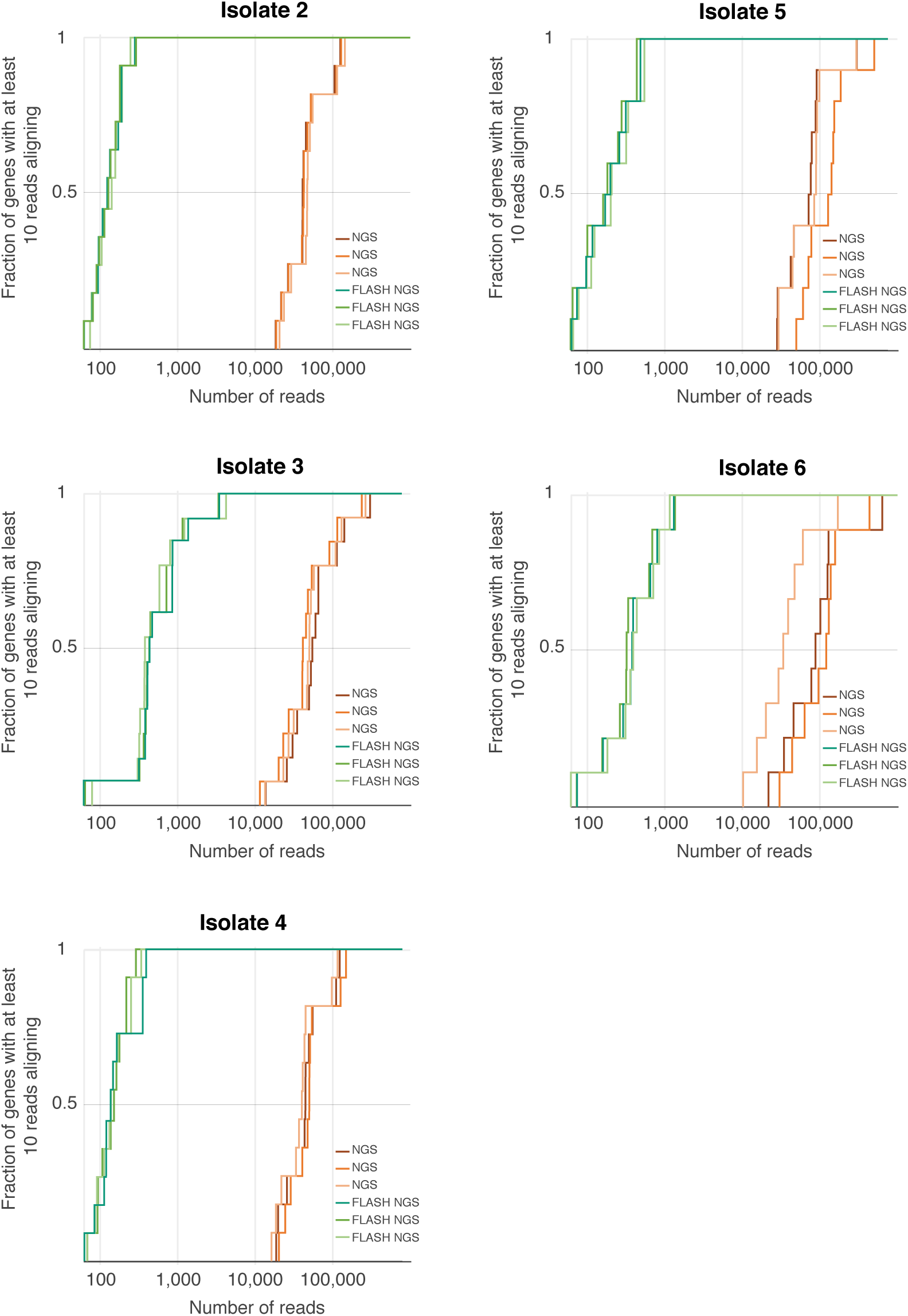
For each FLASH-NGS sample, a sequencing depth between 500 and 5,000 reads was sufficient to recover 10 or more reads per gene for 100% of targeted genes. For NGS alone, at least 100-fold higher sequencing depth was required to achieve this threshold.

**Figure S3.**
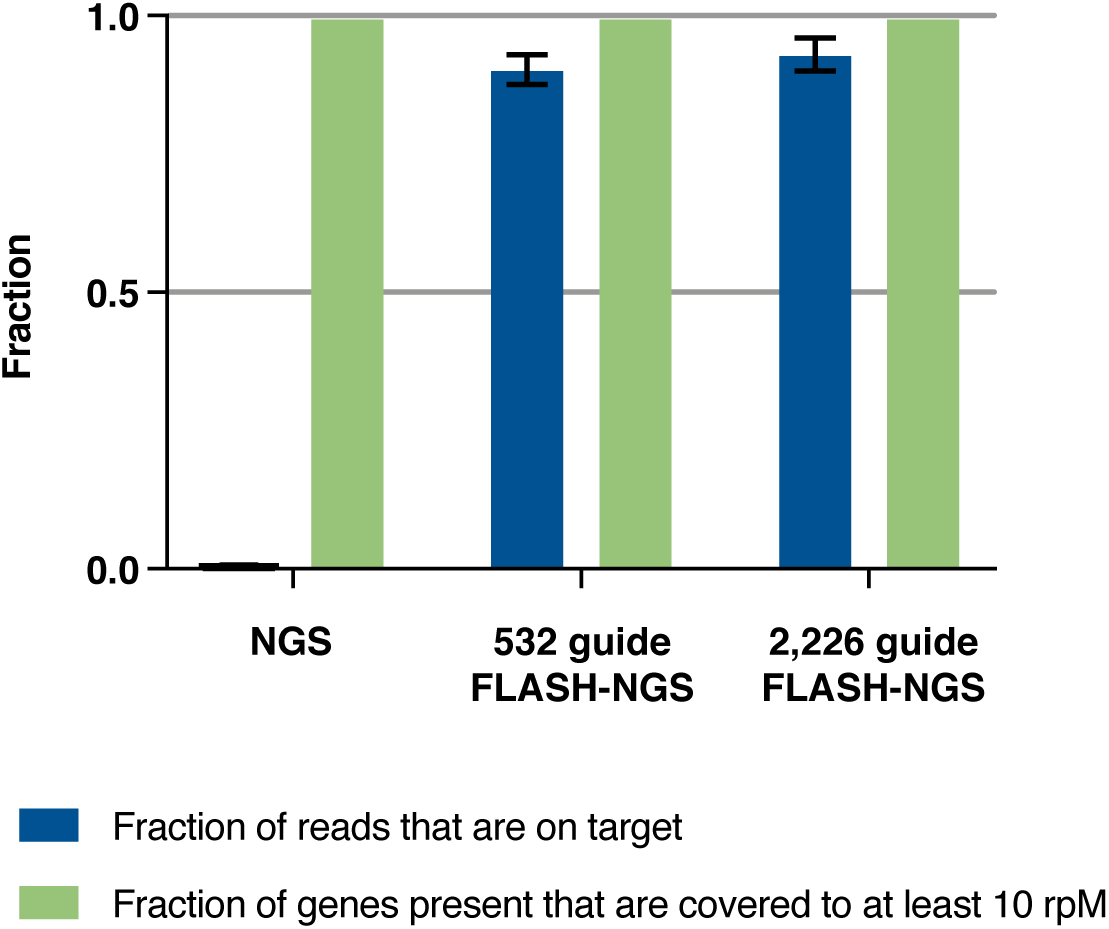
An extended set of 2,226 AMR-targeted guide RNAs was constructed and used in FLASH-NGS of isolate 1. Reads were filtered and aligned to the extended set of genes. One additional gene not targeted in the pilot guide RNA, blaZ-35, was detected and enriched with FLASH (see Table S5).

**Figure S4.**
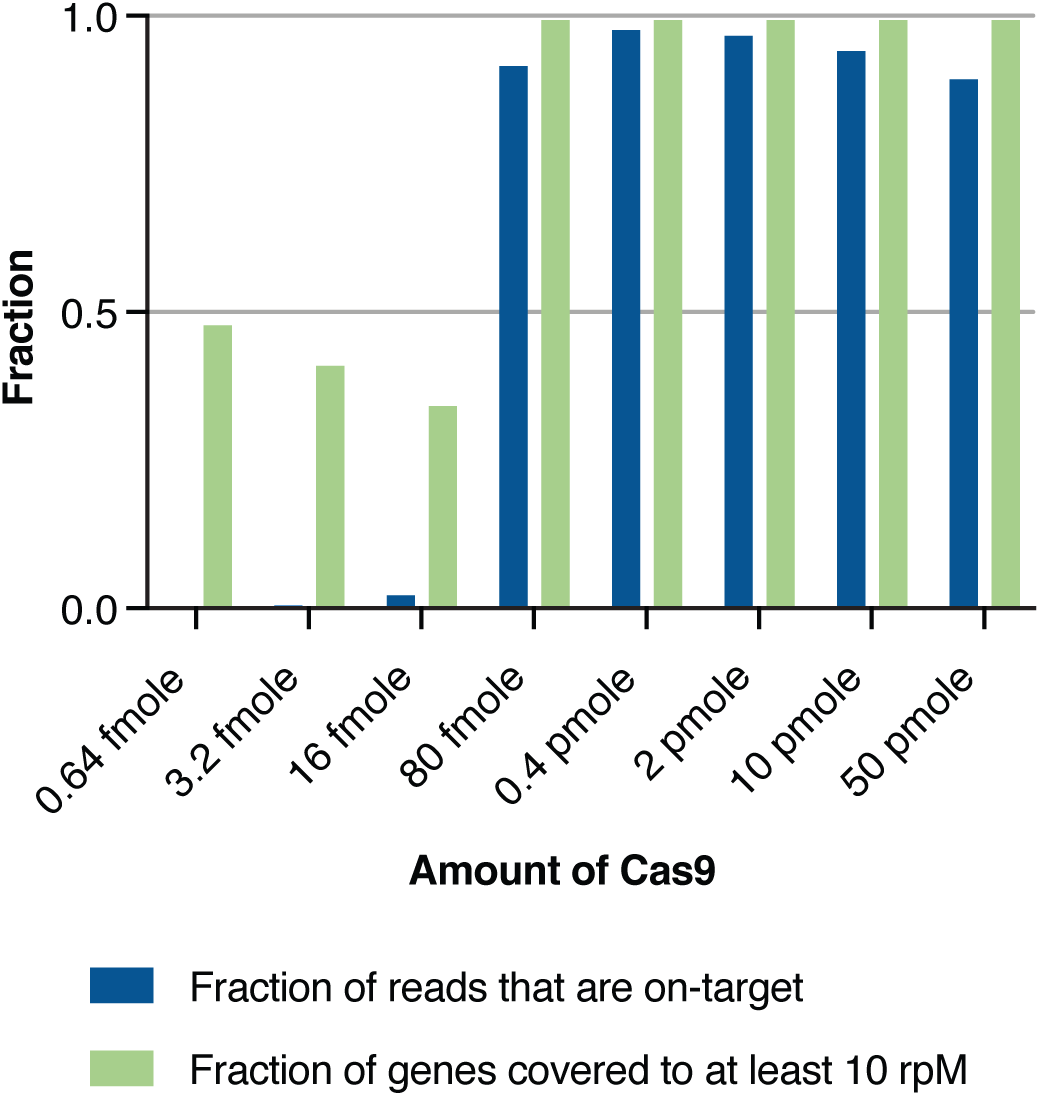
Varying amounts of Cas9 was used in FLASH-NGS of isolate 1. Each data point represents a single replicate.

**Figure S5.**
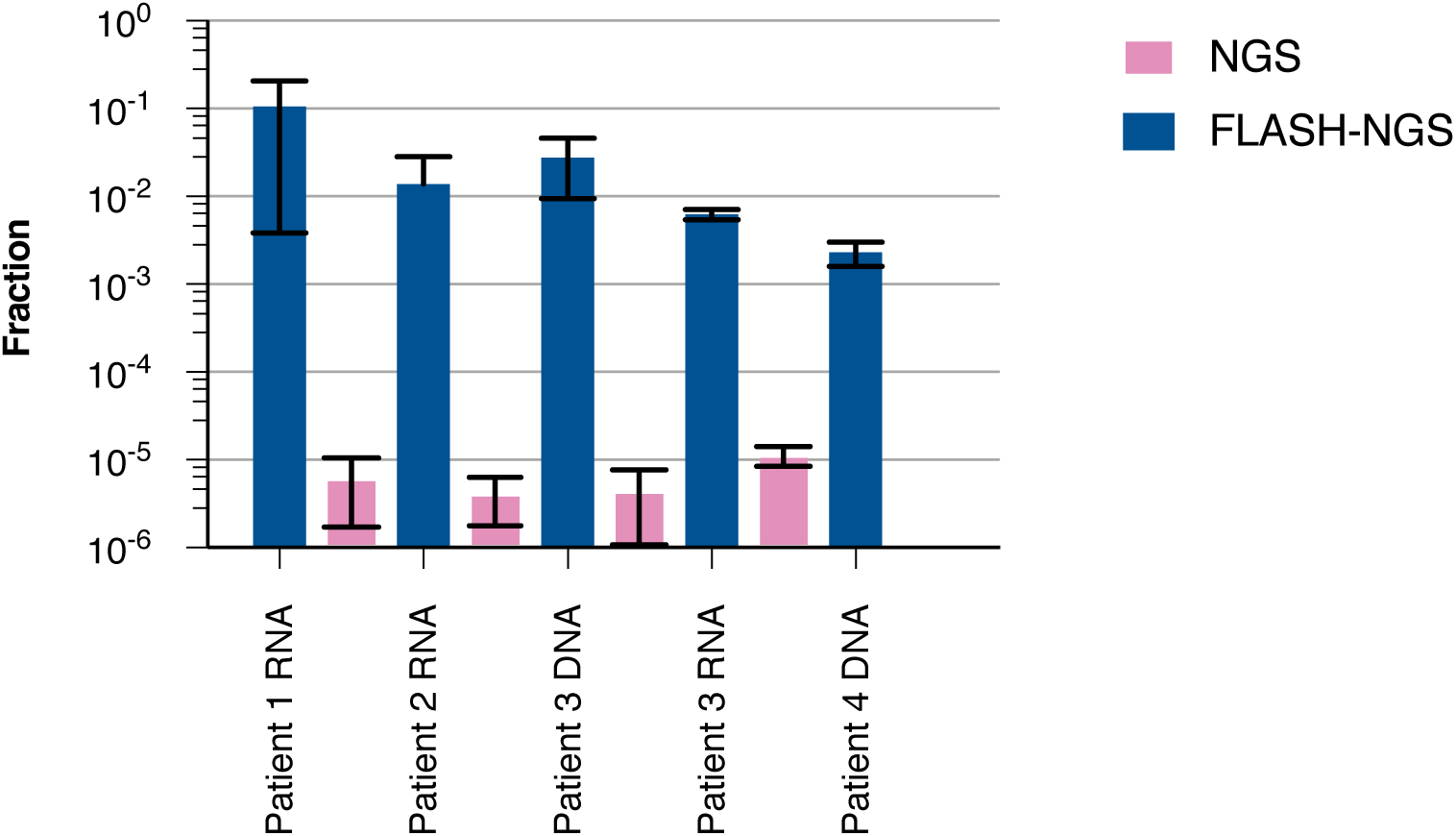
Fraction of on-target reads in NGS and FLASH-NGS experiments on metagenomic patient samples.

**Figure S6.**
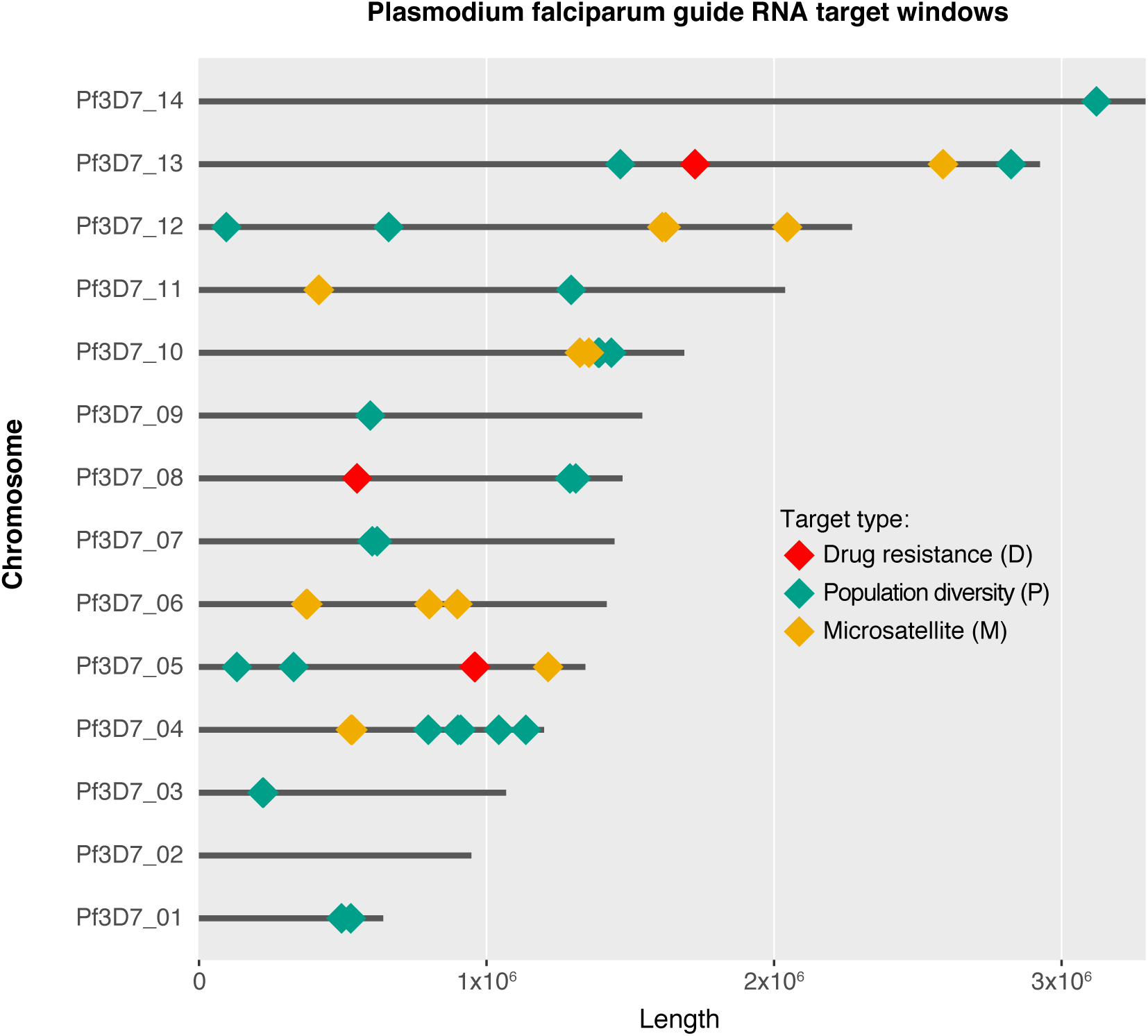
Forty-eight *P. falciparum* genomic loci were targeted with FLASH-NGS.

**Figure S7.**
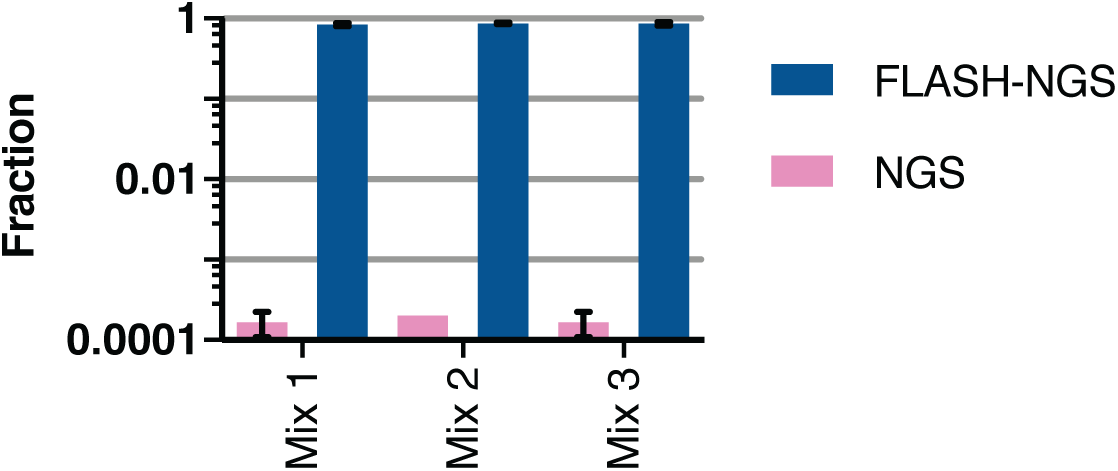
Fraction of on-target reads in NGS and FLASH-NGS experiments on *P. falciparum* lab strain mixture samples.

**Figure S8.**
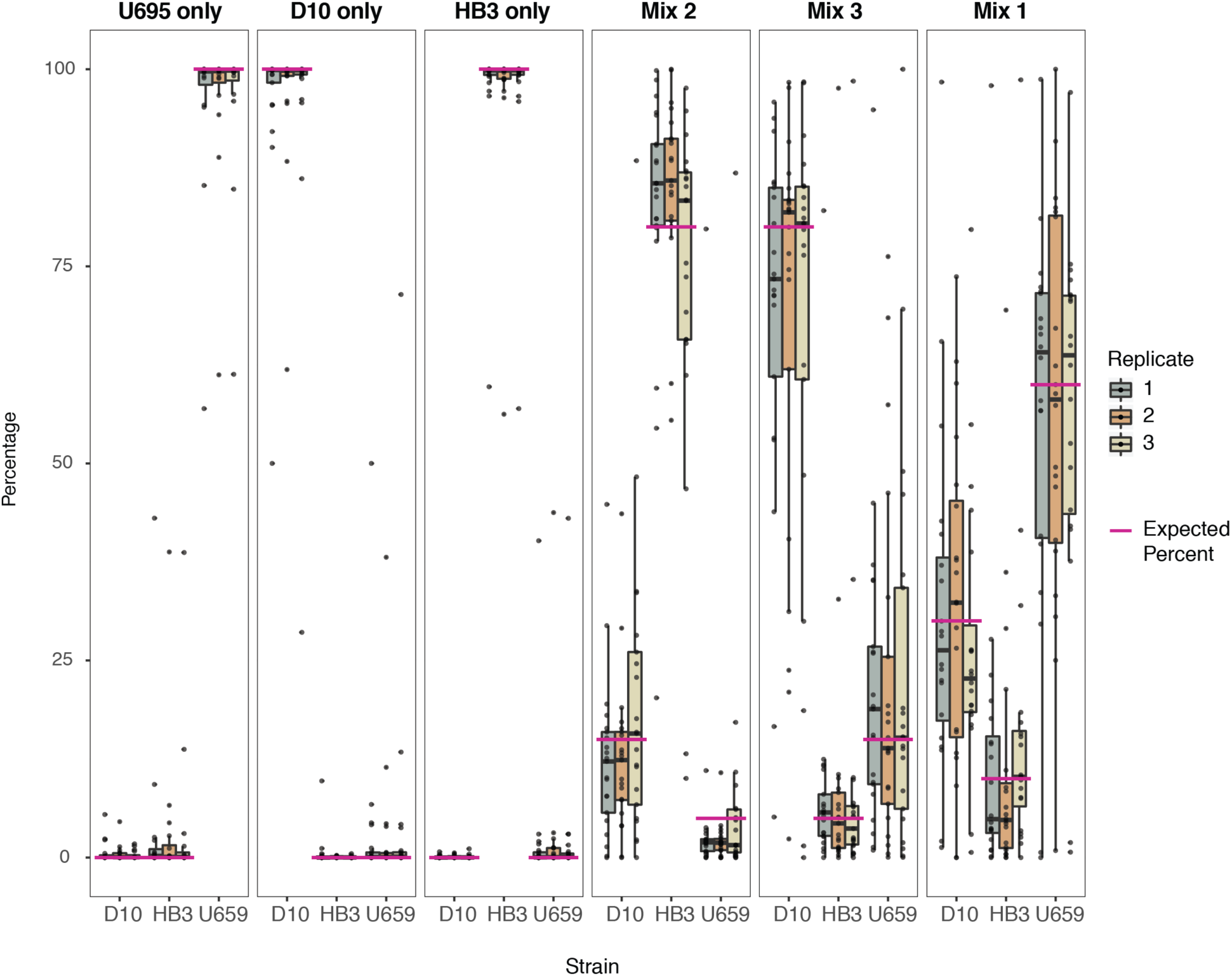
Three lab strains and three lab strain mixtures (see Fig. 4) were subjected to FLASH-NGS. For 21 of the 48 target windows, haplotype sequences uniquely identify each strain. Thus, for those windows, the observed percentages of reads belonging to each strain are plotted. Dots represent individual target windows. Boxes represent median and interquartile range. Pink lines indicate expected strain percentages. Windows included are: P-01, P-03, P-05, P-06, P-07, P-08, P-11, P-12, P-13, P-14, P-15, P-17, P-18, P-21, P-23, P-24, P-25, M-04, M-12, M-15, and M-17.

**Figure S9.**
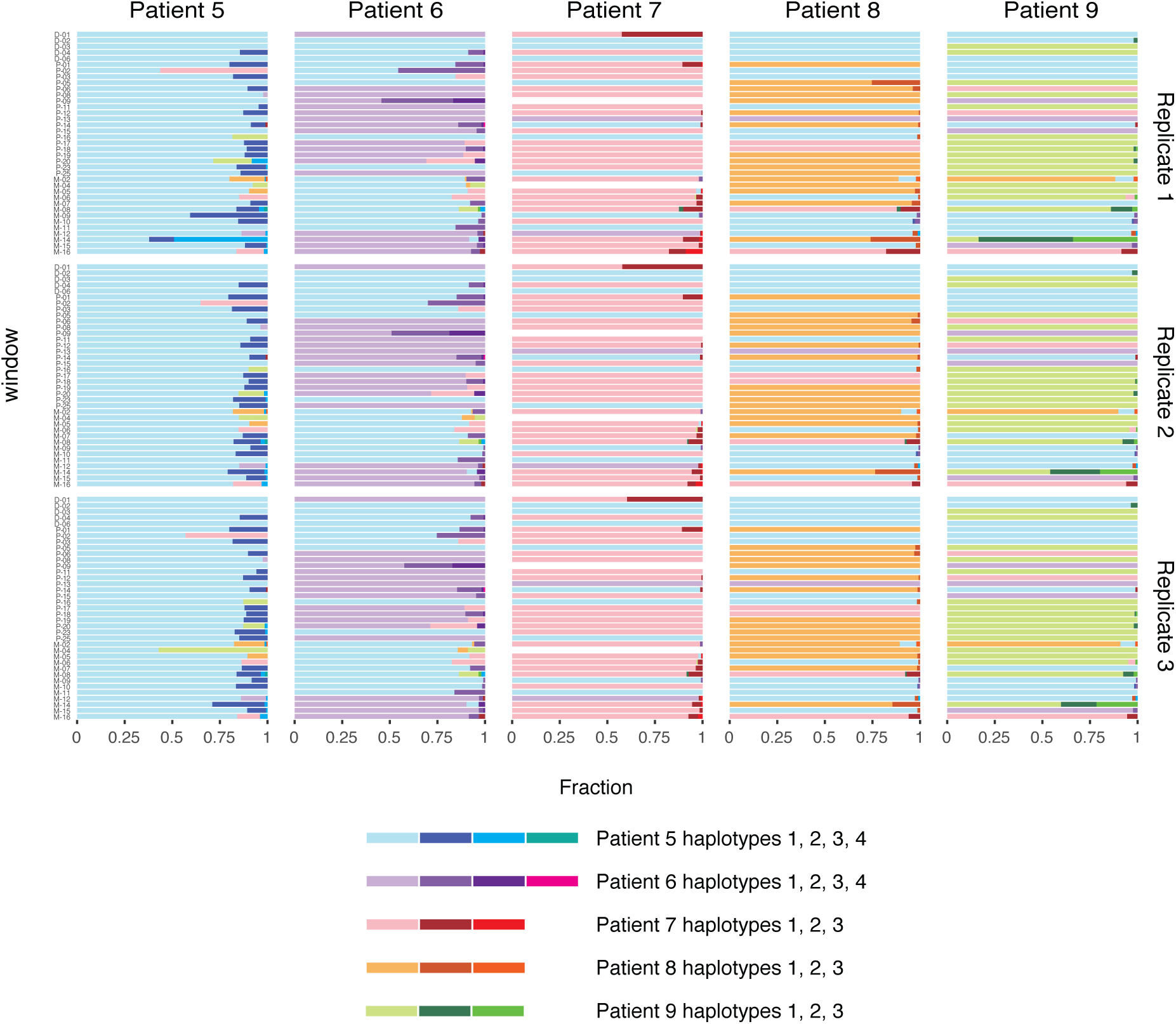
SeekDeep output indicating varying haplotype sequences for five patients infected with *P. falciparum*. Patients 5 and 6 have up to four haplotypes represented in each window; patients 7, 8 and 9 have up to three. Some windows had identical sequences across multiple patients, but none of these patients shared a complete set of identical primary haplotype sequences, suggesting that five different strains accounted for the primary infections in these five individuals. Haplotypes were colored as follows: First, primary haplotype sequences in patient 5, plus any identical sequences in the other patients, were colored light blue. Then primary haplotype sequences in patient 6, plus any identical sequences in other patients, were colored light purple. This was repeated for patients 7 (pink), 8 (light orange) and nine (light green). Next, secondary haplotype sequences in patient 5, plus any identical sequences in the other patients, were colored dark blue. This was repeated for secondary haplotypes in the remaining patients, then for tertiary haplotypes, and finally for quaternary haplotypes.

**Table S6.** Phenotypic drug resistance data for bacterial isolates involved in this study.

| Antibiotic | S. aureus isolates |  |  |  |  |  |
| --- | --- | --- | --- | --- | --- | --- |
|  | Isolate 1 | Isolate 2 | Isolate 3 | Isolate 4 | Isolate 5 | Isolate 6 |
| Clindamycin | R **ermC** | R **unknown** | R **ermC** | R **unknown** | R **ermC** | R **unknown** |
| Nafcillin | R **mecA** | R **mecA** | R **mecA** | R **mecA** | S | S |
| TMP/SMX | S | S | R **dfrG** | S | S | n/a |
| Ciprofloxacin | S | R **gyrA S84L** | S | R **unknown** | S | S |
| Vancomycin | S | S | S | S | S | S |

|  | Strains isolated from patient samples |  |  |  |
| --- | --- | --- | --- | --- |
|  | *S. aureus* |  | *E. faecium* |  |
|  | Patient 1 | Patient 2* | Patient 3 | Patient 4 |
| Clindamycin | S | R **ermA, ermC** | n/a **ermB** | n/a **ermB** |
| Nafcillin | R **mecA** | R **mecA** | n/a | n/a |
| TMP/SMX | S | R **dfrC** | n/a **dfrF, sul1** | n/a **dfrG** |
| Ciprofloxacin | S | R **S. aureus gyrA S84L** | n/a | n/a |
| Vancomycin | S | S | R **vanA** | R **vanA** |
| Rifampicin | n/a | n/a **S. aureus rpoB H481Y** | n/a | n/a |

R: resistant
S: sensitive
n/a: not tested
**blue**: detected with FLASH
\* Note: Patient 2 was infected with Staphylococcus, Citrobacter and Pseudomonas. ErmA, ermC and dfrC are not known to be present in Citrobacter or Pseudomonas and are most likely derived from *S. aureus* in this patient.

## REFERENCES

1. O’Neill, J. (2016) Tackling Drug-Resistant Infections Globally: Final Report and Recommendations The Review on Antimicrobial Resistance.

2. Didelot, X., Bowden, R., Wilson, D.J., Peto, T.E.A. and Crook, D.W. (2012) Transforming clinical microbiology with bacterial genome sequencing. Nat. Rev. Genet., 13, 601–612.

3. Zaas, A.K., Garner, B.H., Tsalik, E.L., Burke, T., Woods, C.W. and Ginsburg, G.S. (2014) The current epidemiology and clinical decisions surrounding acute respiratory infections. Trends Mol Med, 20, 579–588.

4. van der Eerden, M.M., Vlaspolder, F., de Graaff, C.S., Groot, T., Bronsveld, W., Jansen, H.M. and Boersma, W.G. (2005) Comparison between pathogen directed antibiotic treatment and empirical broad spectrum antibiotic treatment in patients with community acquired pneumonia: a prospective randomised study. Thorax, 60, 672–678.

5. Amato, R., Pearson, R.D., Almagro-Garcia, J., Amaratunga, C., Lim, P., Suon, S., Sreng, S., Drury, E., Stalker, J., Miotto, O., et al. (2018) Origins of the current outbreak of multidrug-resistant malaria in southeast Asia: a retrospective genetic study. The Lancet Infectious Diseases, 18, 337–345.

6. Rosenthal, P.J. (2013) The interplay between drug resistance and fitness in malaria parasites. Mol. Microbiol., 89, 1025–1038.

7. Nsanzabana, C., Djalle, D., Guérin, P.J., Ménard, D. and González, I.J. (2018) Tools for surveillance of anti-malarial drug resistance: an assessment of the current landscape. Malaria Journal, 17, 75.

8. Greenhouse, B., Dokomajilar, C., Hubbard, A., Rosenthal, P.J. and Dorsey, G. (2007) Impact of Transmission Intensity on the Accuracy of Genotyping To Distinguish Recrudescence from New Infection in Antimalarial Clinical Trials. Antimicrob. Agents Chemother., 51, 3096–3103.

9. Yozwiak, N.L., Skewes-Cox, P., Stenglein, M.D., Balmaseda, A., Harris, E. and DeRisi, J.L. (2012) Virus Identification in Unknown Tropical Febrile Illness Cases Using Deep Sequencing. PLoS Neglected Tropical Diseases, 6, e1485.

10. Wilson, M.R., Naccache, S.N., Samayoa, E., Biagtan, M., Bashir, H. and Yu, G. (2014) Actionable diagnosis of neuroleptospirosis by next-generation sequencing. N Engl J Med., 370.

11. Langelier, C., Zinter, M.S., Kalantar, K., Yanik, G.A., Christenson, S., O’Donovan, B., White, C., Wilson, M., Sapru, A., Dvorak, C.C., et al. (2017) Metagenomic Sequencing Detects Respiratory Pathogens in Hematopoietic Cellular Transplant Patients. Am J Respir Crit Care Med, 197, 524–528.

12. Urbaniak, C., Sielaff, A.C., Frey, K.G., Allen, J.E., Singh, N., Jaing, C., Wheeler, K. and Venkateswaran, K. (2018) Detection of antimicrobial resistance genes associated with the International Space Station environmental surfaces. Scientific Reports, 8.

13. Jinek, M., Chylinski, K., Fonfara, I., Hauer, M., Doudna, J.A. and Charpentier, E. (2012) A programmable dual-RNA-guided DNA endonuclease in adaptive bacterial immunity. Science., 337.

14. Gootenberg, J.S., Abudayyeh, O.O., Lee, J.W., Essletzbichler, P., Dy, A.J., Joung, J., Verdine, V., Donghia, N., Daringer, N.M., Freije, C.A., et al. (2017) Nucleic acid detection with CRISPR-Cas13a/C2c2. Science, 356, 438.

15. Gootenberg, J.S., Abudayyeh, O.O., Kellner, M.J., Joung, J., Collins, J.J. and Zhang, F. (2018) Multiplexed and portable nucleic acid detection platform with Cas13, Cas12a, and Csm6. Science, 10.1126/science.aaq0179.

16. East-Seletsky, A., O’Connell, M.R., Knight, S.C., Burstein, D., Cate, J.H.D., Tjian, R. and Doudna, J.A. (2016) Two distinct RNase activities of CRISPR-C2c2 enable guide-RNA processing and RNA detection. Nature, 538, 270–273.

17. Chen, J.S., Ma, E., Harrington, L.B., Da Costa, M., Tian, X., Palefsky, J.M. and Doudna, J.A. (2018) CRISPR-Cas12a target binding unleashes indiscriminate single-stranded DNase activity. Science, 360, 436.

18. Gu, W., Crawford, E.D., O’Donovan, B.D., Wilson, M.R., Chow, E.D., Retallack, H. and DeRisi, J.L. (2016) Depletion of Abundant Sequences by Hybridization (DASH): using Cas9 to remove unwanted high-abundance species in sequencing libraries and molecular counting applications. Genome Biology, 17, 1–13.

19. Jia, B., Raphenya, A.R., Alcock, B., Waglechner, N., Guo, P., Tsang, K.K., Lago, B.A., Dave, B.M., Pereira, S., Sharma, A.N., et al. (2017) CARD 2017: expansion and model-centric curation of the comprehensive antibiotic resistance database. Nucleic Acids Research, 45, D566–D573.

20. Zankari, E., Hasman, H., Cosentino, S., Vestergaard, M., Rasmussen, S., Lund, O., Aarestrup, F.M. and Larsen, M.V. (2012) Identification of acquired antimicrobial resistance genes. Journal of Antimicrobial Chemotherapy, 67, 2640–2644.

21. Ruby, J.G., Bellare, P. and DeRisi, J.L. (2013) PRICE: software for the targeted assembly of components of (meta) genomic sequence data. G3 Genes Genomes Genetics, 3.

22. Langmead, B. and Salzberg, S.L. (2012) Fast gapped-read alignment with Bowtie 2. Nat Methods., 9.

23. Oyola, S.O., Ariani, C.V., Hamilton, W.L., Kekre, M., Amenga-Etego, L.N., Ghansah, A., Rutledge, G.G., Redmond, S., Manske, M., Jyothi, D., et al. (2016) Whole genome sequencing of Plasmodium falciparum from dried blood spots using selective whole genome amplification. Malaria Journal, 15, 597.

24. Aurrecoechea, C., Brestelli, J., Brunk, B.P., Dommer, J., Fischer, S., Gajria, B., Gao, X., Gingle, A., Grant, G., Harb, O.S., et al. (2009) PlasmoDB: a functional genomic database for malaria parasites. Nucleic Acids Res., 37, D539-543.

25. Li, H., Handsaker, B., Wysoker, A., Fennell, T., Ruan, J. and Homer, N. (2009) The Sequence Alignment/Map format and SAMtools. Bioinforma Oxf Engl., 25.

26. Quinlan - 2014 - BEDTools The Swiss-Army Tool for Genome Feature A.pdf.

27. Haeussler, M., Schönig, K., Eckert, H., Eschstruth, A., Mianné, J., Renaud, J.-B., Schneider-Maunoury, S., Shkumatava, A., Teboul, L., Kent, J., et al. (2016) Evaluation of off-target and on-target scoring algorithms and integration into the guide RNA selection tool CRISPOR. Genome Biology, 17.

28. Listgarten, J., Weinstein, M., Kleinstiver, B.P., Sousa, A.A., Joung, J.K., Crawford, J., Gao, K., Hoang, L., Elibol, M., Doench, J.G., et al. (2018) Prediction of off-target activities for the end-to-end design of CRISPR guide RNAs. Nature Biomedical Engineering, 2, 38.

29. O’Neill, A.J., Huovinen, T., Fishwick, C.W.G. and Chopra, I. (2006) Molecular Genetic and Structural Modeling Studies of Staphylococcus aureus RNA Polymerase and the Fitness of Rifampin Resistance Genotypes in Relation to Clinical Prevalence. Antimicrobial Agents and Chemotherapy, 50, 298–309.

30. Fairhurst, R.M. and Dondorp, A.M. (2016) Artemisinin-Resistant Plasmodium falciparum Malaria. In Scheld, W.M., Hughes, J.M., Whitley, R.J. (eds), Emerging infections 10. American Society of Microbiology, pp. 409–429.

31. Pearce, R.J., Pota, H., Evehe, M.-S.B., Bâ, E.-H., Mombo-Ngoma, G., Malisa, A.L., Ord, R., Inojosa, W., Matondo, A., Diallo, D.A., et al. (2009) Multiple Origins and Regional Dispersal of Resistant dhps in African Plasmodium falciparum Malaria. PLoS Medicine, 6, e1000055.

32. Hathaway, N.J., Parobek, C.M., Juliano, J.J. and Bailey, J.A. (2018) SeekDeep: single-base resolution de novo clustering for amplicon deep sequencing. Nucleic Acids Res, 46, e21–e21.

## REFERENCES

1. Langelier, C., Kalantar, K.L., Moazed, F., Wilson, M.R., Crawford, E., Deiss, T., Belzer, A., Bolourchi, S., Caldera, S., Fung, M., et al. (2018) Integrating Host Response and Unbiased Microbe Detection for Lower Respiratory Tract Infection Diagnosis in Critically Ill Adults. 10.1101/341149.

2. Plowe, C.V., Djimde, A., Bouare, M., Doumbo, O. and Wellems, T.E. (1995) Pyrimethamine and Proguanil Resistance-Conferring Mutations in Plasmodium falciparum Dihydrofolate Reductase: Polymerase Chain Reaction Methods for Surveillance in Africa. The American Journal of Tropical Medicine and Hygiene, 52, 565–568.

3. Hofmann, N., Mwingira, F., Shekalaghe, S., Robinson, L.J., Mueller, I. and Felger, I. (2015) Ultra-Sensitive Detection of Plasmodium falciparum by Amplification of Multi-Copy Subtelomeric Targets. PLOS Medicine, 12, e1001788.

4. Sundararaman, S.A., Plenderleith, L.J., Liu, W., Loy, D.E., Learn, G.H., Li, Y., Shaw, K.S., Ayouba, A., Peeters, M., Speede, S., et al. (2016) Genomes of cryptic chimpanzee Plasmodium species reveal key evolutionary events leading to human malaria. Nature Communications, 7, 11078.

5. Oyola, S.O., Ariani, C.V., Hamilton, W.L., Kekre, M., Amenga-Etego, L.N., Ghansah, A., Rutledge, G.G., Redmond, S., Manske, M., Jyothi, D., et al. (2016) Whole genome sequencing of Plasmodium falciparum from dried blood spots using selective whole genome amplification. Malaria Journal, 15, 597.

6. Gu, W., Crawford, E.D., O’Donovan, B.D., Wilson, M.R., Chow, E.D., Retallack, H. and DeRisi, J.L. (2016) Depletion of Abundant Sequences by Hybridization (DASH): using Cas9 to remove unwanted high-abundance species in sequencing libraries and molecular counting applications. Genome Biology, 17, 1–13.

7. Wu, X., Scott, D.A., Kriz, A.J., Chiu, A.C., Hsu, P.D., Dadon, D.B., Cheng, A.W., Trevino, A.E., Konermann, S., Chen, S., et al. (2014) Genome-wide binding of the CRISPR endonuclease Cas9 in mammalian cells. Nature Biotechnology, 32, 670–676.

8. Boyle, E.A., Andreasson, J.O.L., Chircus, L.M., Sternberg, S.H., Wu, M.J., Guegler, C.K., Doudna, J.A. and Greenleaf, W.J. (2017) High-throughput biochemical profiling reveals sequence determinants of dCas9 off-target binding and unbinding. Proceedings of the National Academy of Sciences, 114, 5461–5466.

9. Gurobi Optimization, LLC (2018) Gurobi Optimizer Reference Manual.

10. Ruby, J.G., Bellare, P. and DeRisi, J.L. (2013) PRICE: software for the targeted assembly of components of (meta) genomic sequence data. G3 Genes Genomes Genetics, 3.

11. Li, H., Handsaker, B., Wysoker, A., Fennell, T., Ruan, J. and Homer, N. (2009) The Sequence Alignment/Map format and SAMtools. Bioinforma Oxf Engl., 25.

12. Kearse, M., Moir, R., Wilson, A., Stones-Havas, S., Cheung, M., Sturrock, S., Buxton, S., Cooper, A., Markowitz, S., Duran, C., et al. (2012) Geneious Basic: an integrated and extendable desktop software platform for the organization and analysis of sequence data. Bioinformatics, 28, 1647–1649.

13. Aurrecoechea, C., Brestelli, J., Brunk, B.P., Dommer, J., Fischer, S., Gajria, B., Gao, X., Gingle, A., Grant, G., Harb, O.S., et al. (2009) PlasmoDB: a functional genomic database for malaria parasites. Nucleic Acids Res., 37, D539-543.

14. Quinlan - 2014 - BEDTools The Swiss-Army Tool for Genome Feature A.pdf.

15. Hathaway, N.J., Parobek, C.M., Juliano, J.J. and Bailey, J.A. (2018) SeekDeep: single-base resolution de novo clustering for amplicon deep sequencing. Nucleic Acids Res, 46, e21–e21.

